# Mucosal immune stimulation with HSV-2 and polyICLC boosts control of viremia in SIVΔNef vaccinated rhesus macaques with breakthrough SIV infection

**DOI:** 10.1101/2020.06.02.129494

**Authors:** Meropi Aravantinou, Olga Mizenina, Thilo Brill, Jessica Kenney, Christine Timmons, Ines Frank, Agegnehu Gettie, Brooke Grasperge, James Blanchard, Andres Salazar, Jeffrey D. Lifson, Melissa Robbiani, Nina Derby

**Author notes:** Corresponding author, Nina Derby, Population Council, 1230 York Avenue, New York, NY, 10065.

## Abstract

Development of an effective human immunodeficiency virus (HIV) vaccine is among the highest priorities in the biomedical research agenda. Adjuvants enhance vaccine efficacy, but in the case of HIV, strong or inappropriate immune activation may undermine protection by increasing HIV susceptibility. Co-infection with immunomodulatory pathogens may also impact vaccine efficacy. In the rhesus macaque rectal SIVΔNef live attenuated vaccine model, we utilized a low virulence HSV-2 infection and the double-stranded RNA viral mimic polyICLC as tools to probe the effects of distinct types of immune activation on HIV vaccine efficacy and explore novel correlates of protection from wild type SIV. Rectally administered HSV-2 and polyICLC impacted the protection conferred by mucosal SIVΔNef vaccination by favoring partial protection in animals with breakthrough infection following virulent SIV challenge (“Controllers”). However, SIVΔNef persistence in blood and tissues did not predict protection in this rectal immunization and challenge model. Non-controllers had similar SIVΔNef viremia as completely protected macaques, and while they tended to have less replication competent SIVΔNef in lymph nodes, controllers had no recoverable virus in the lymph nodes. Non-controllers differed from protected macaques immunologically by having a greater frequency of pro-inflammatory CXCR3^+^CCR6^+^ CD4 T cells in blood and a monofunctional IFNγ-dominant CD8 T cell response in lymph nodes. Controller phenotype was associated with heightened IFNα production during acute SIV infection and a greater frequency of CXCR5^+^ CD4 T cells in blood pre-challenge despite a lower frequency of cells with the T follicular helper (Tfh) cell phenotype in blood and lymph nodes. Our results establish novel correlates of immunological control of SIV infection while reinforcing the potential importance of T cell functionality and location in SIVΔNef efficacy. Moreover, this work highlights that triggering of mucosal immunity can aid mucosal vaccine strategies rather than undermine protection.

**AUTHOR SUMMARY:** An efficacious HIV vaccine is essential to contain the HIV pandemic. Vaccine-mediated protection from HIV may be either enhanced or obstructed by mucosal immune activation; thus, the impact of adjuvants and underlying co-infections that lead to immune activation needs to be evaluated. Using the SIV macaque model, we set out to study the impact of underlying infection with HSV-2 or treatment with the adjuvant polyICLC on rectal immunization with the live attenuated vaccine SIVΔNef. We found that neither stimulus impacted complete protection from SIV; however, the combination of HSV-2 and polyICLC improved control of infection in animals that were not completely protected. Compared with non-controller macaques, controllers had less inflammatory T cells before SIV challenge as well as greater gene expression of IFNα and more functional SIV-specific T cells after infection. The results add to our understanding of the mechanisms of SIVΔNef protection and demonstrate that mucosal immune activation does not necessarily undermine protection in mucosal vaccination against HIV.

## INTRODUCTION

Designing an efficacious HIV vaccine is a difficult task by any measure due to the high level of viral diversity, immune escape, and early seeding of cellular and anatomical viral reservoirs. Any efforts that can reduce the barriers to efficacy should be considered. Mucosal vaccines offer advantages over parenteral vaccines in their enhanced ability to position antigen-specific immune responses where they are needed to curb sexual transmission (1). However, vaccine-associated immune activation in the mucosa, including that provided by the adjuvants employed to augment and focus the immune response and by underlying co-infections, may have the undesired side effect of increasing available target cells for HIV infection. When lack of efficacy and enhancement of HIV acquisition were found in large-scale clinical trials of Adenovirus (Ad)- based HIV vaccines (2, 3), a link with mucosal T cell activation was hypothesized (4, 5). Given the potential dichotomy between appropriate and pathological mucosal immune activation, it is critical to tease out the impacts of mucosal stimuli on HIV vaccine efficacy.

As a model system to address the question, we used the SIVmac239ΔNef (SIVΔNef) live attenuated vaccine (LAV). SIVΔNef is among the best characterized SIV vaccines in macaques and a benchmark for the potency of protective immunity that should be elicited by an efficacious (but safer) HIV vaccine in humans. SIVΔNef engages multiple arms of the immune system, including humoral and T cell-mediated responses and even mucosal immunity in response to systemic vaccination (6–9). Thus, it represents an excellent choice for examining the effect of immune stimuli on different components of protection. While a number of studies have found that high-dose intravenous SIVΔNef immunization completely protects macaques against intravenous or mucosal SIV challenge (6–14), we have shown that mucosal immunization with a lower dose of SIVΔNef via the rectal mucosa is less efficient in blocking rectal SIV transmission (15). The presence of breakthrough infections within an immunized cohort allows for the examination of factors that improve or disrupt immunity. Throughout the large body of literature on immunization of macaques with LAVs including SIVΔNef, a number of correlates of vaccine protection have been identified, including persistence of the LAV (6), especially in the PD-1^+^CXCR5^+^CD200^+^ T follicular helper (Tfh) cell subset of CD4 T cells that reside within the lymph node B cell follicles (16). This antigen persistence is thought to drive the production of the effector differentiated SIV-specific T cells (6, 8) and antibodies (6, 7, 9) that have been correlated with protection.

To explore the impact of mucosal stimulation on rectal mucosa SIVΔNef immunization and challenge in macaques (15, 17), we selected polyICLC and herpes simplex virus-2 (HSV-2) as model stimuli with divergent mechanisms of cell activation and potentially divergent immunological effects. PolyICLC is an adjuvant that activates dendritic cells (DCs) and T cells (15, 18–20), and we found that rectal administration of polyICLC just prior to SIVΔNef improves protection against SIV challenge (15). Infection with HSV-2 in humans confers a 3-fold enhancement in HIV infection risk (21) through mechanisms including mucosal inflammation, upregulation of HIV receptors, and recruitment and persistence of HIV target cells in the genital and anorectal mucosa (17, 22–28). HSV-2 infection in rhesus macaques recapitulates many of the features of infection in humans but is less pathogenic with a more restricted pattern of shedding (22, 29–31). Nonetheless, high dose vaginal and rectal HSV-2 inoculation in macaques creates a pro-inflammatory state with T cell activation in mucosa and blood (23, 24) that mirrors similar findings in humans (28). Moreover, in opposition to the effect of polyICLC, post-vaccination acute rectal HSV-2 infection in macaques dampened the protective efficacy of SIVΔNef in association with increased mucosal inflammation (17).

In the current study, we utilized a repeated low dose HSV-2 infection model together with polyICLC to generate a matrix of differing types of immune activation and differing associated levels of protection that could facilitate understanding how immune activation impacts protection by SIVΔNef and possibly uncover novel correlates of protection. Testing of how an underlying low virulence rectal HSV-2 infection impacts SIVΔNef efficacy in macaques might also inform on the impact of subclinical human HSV-2 infection on mucosal HIV vaccines.

We found that both HSV-2 infection and polyICLC increased acquisition of SIVΔNef (vaccine take) following a low dose intrarectal inoculation. Notably, in animals in which there was a take of the SIVΔNef, neither HSV-2 nor polyICLC pre-treatment impacted the elicitation of complete protection by SIVΔNef. However, the combination of both mucosal stimuli boosted the control of SIV in animals with breakthrough infection, revealing a novel partial protection “controller” phenotype. In this model, control (vs. non-control) was not directly predicted by SIVΔNef persistence but rather by increased IFNα production during acute SIV infection as well as a heightened frequency of CXCR5^high^ CD4 T cells in blood, despite a lower frequency of such cells in lymph nodes and a lower frequency of Tfh cells. Controllers also exhibited lower expression of CD40L on their SIV-specific CD8 T cells than completely protected macaques. Moreover, we found a monofunctional CD8 T cell response in non-controllers that was not seen in controllers or completely protected macaques. Our results reinforce the importance of T cell functionality and tissue localization, as well as an aptly timed innate response in control of SIV infection. This provides a platform for future testing of the importance of immune modifying conditions on HIV vaccine outcomes.

## RESULTS

### Rectal HSV-2 infection and polyICLC treatment promote an SIV controller phenotype in breakthrough infections in SIVΔNef mucosal vaccinees

We evaluated the impact of two mucosal stimuli – HSV-2 and polyICLC – on rectal SIVΔNef-mediated protection from SIVmac239 in a cohort of 31 rhesus macaques. The macaques were divided in 4 treatment groups: control (n=10), HSV-2 (n=11), polyICLC (n=4), and HSV-2/polyICLC (n=6) (**Fig. 1A**). We hypothesized that HSV-2 would exert an immune activating effect, undermining SIVΔNef-mediated protection and that polyICLC would boost protection and possibly buffer dysregulating effects of HSV-2. A 10-week regimen of twice-weekly mucosal treatments with 10^7^ pfu of HSV-2 and/or 1mg of polyICLC was employed. This regimen was used to mimic physiological conditions, which may include accumulated effects of repeated exposure to immunomodulatory agents such as HSV-2. In addition, we previously found that repeated vaginal exposure to 10^7^ pfu of HSV-2 (together with lentivirus) was associated with sustained HSV-2 shedding in the macaques’ vaginal mucosa (29, 30). Since we aimed to maximize the effects of stimuli and predicted that frequent shedding would more strongly undermine SIVΔNef-mediated protection, we followed a similar challenge protocol herein. Following the treatment regimens, all macaques were exposed rectally to 1×10^3^ TCID_50_ SIVΔNef. In contrast to higher dose mucosal exposures or intravenous inoculation with SIVΔNef, which results in near universal take of SIVΔNef and associated protection, this low dose of the vaccine was utilized in order to create a control group of animals that were exposed to the stimuli and vaccine but did not become infected with SIVΔNef (no vaccine take) and to observe effects of the stimuli on vaccine take (15). As we were unable to exclude or fully balance the inclusion of protective MHC alleles between groups (**Fig. S1**), we present the data concerning SIV protection and pathogenesis for all animals (**Fig. 1–9**) as well as for the subset of animals lacking the Mamu A*01, B*08, and B*17 alleles (MHC censored data in **Supporting Information**).

**Figure 1.**
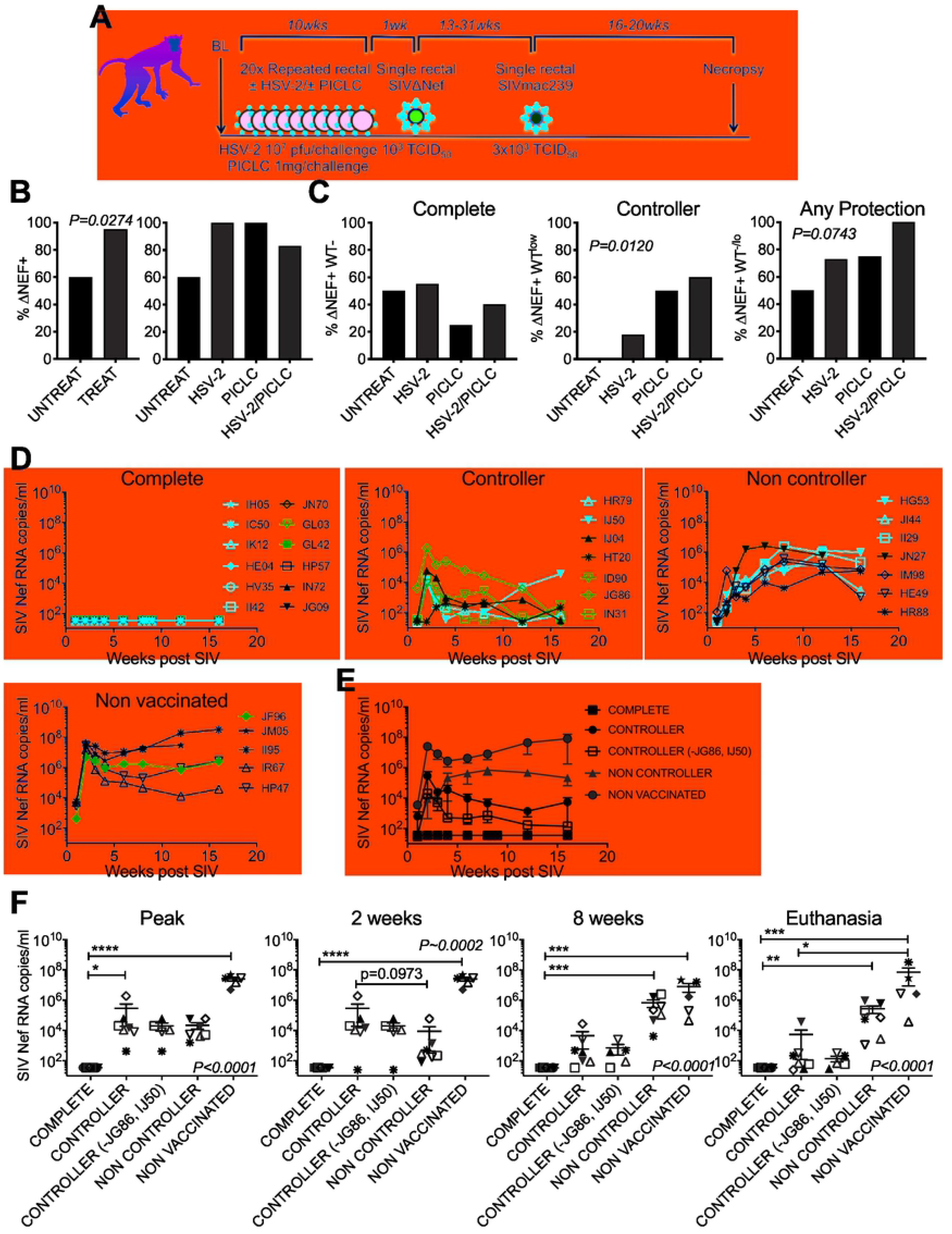
Mucosal immune stimulation promotes SIVΔNef vaccine take and improves protection. **(A)** Schematic of the study design. Thirty-one macaques were divided in 4 treatment groups for 20 treatments in rectal mucosa over 10 weeks. Untreated control group (UNTREAT) animals received PBS (n=10); HSV-2 group (HSV-2) animals received 10^7^ pfu HSV-2 per treatment (n=11); polyICLC group (PICLC) animals received 1 mg polyICLC (Hiltonol®) per treatment (n=4); HSV-2/polyICLC group (HSV-2/PICLC) animals received 10^7^ pfu HSV-2 and 1 mg polyICLC per treatment (n=6). **(B)** Uptake of SIVΔNef. Comparison of all treated (TREAT) vs UNTREAT macaques (left panel) was assessed by Fisher‘s Exact test and comparison between the four treatment groups (right panel) was assessed by Chi Square test for trend. **(C)** Protection from SIV in SIVΔNef+ animals. Complete protection (left panel), controllers (middle panel), and the combination of complete protection and controllers (“Any protection”, right panel) are shown. Comparison of all groups was made by Chi Square test for trend. **(D)** Plasma viral loads for all macaques following SIV challenge according to extent of protection. Macaques that were exposed to but did not become infected with the LAV are designated as “Non Vaccinated”. Treatment groups are indicated by color: HSV-2 (red), PICLC (blue), HSV-2/PICLC (purple), UNTREAT (black). **(E)** Mean ± SEM for the plasma viral load of all animals at each level of protection. **(F)** SIV plasma viremia in different protection groups at distinct phases of SIV infection. Peak viral load indicates the highest viremia between weeks 1 and 4 post-infection. Comparison between groups was made with Kruskal-Wallis test (*P<x*) and Dunns post-test for Kruskal-Wallis *P<0.05*. Approximate Kruskal-Wallis p values are indicated by ~. Dunns p values are *<0.05, **<0.01, ***<0.001.

**Figure 2.**
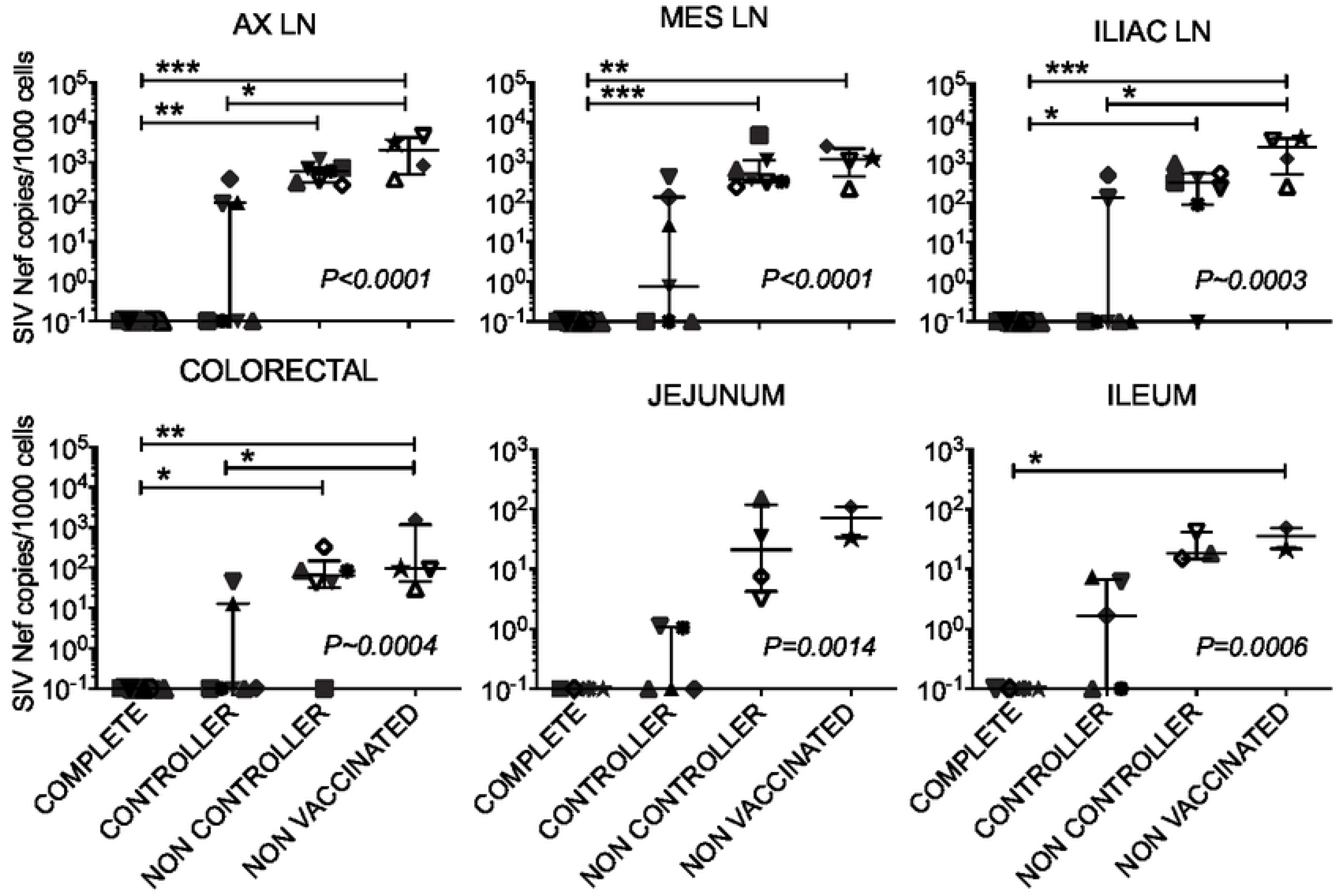
SIV loads in tissues segregate by level of protection. Cell associated SIV *nef* DNA in tissues at necropsy. Tissues examined were axillary (AX), mesenteric (MES), and iliac (ILIAC) lymph nodes (LN) and sections of gut mucosa as shown. Bars and whiskers indicate median ± interquartile range (IQR). SIV copy numbers less than 1 per 10^4^ cells are shown at 1 per 10^4^ cells as the lower limit of detection. Jejunum and ileum were not available from IR67 or HP47. Treatment groups are color coded as in Figure 1. Comparison between groups was made with Kruskal-Wallis test and Dunns post-test as in Figure 1.

**Figure 3.**
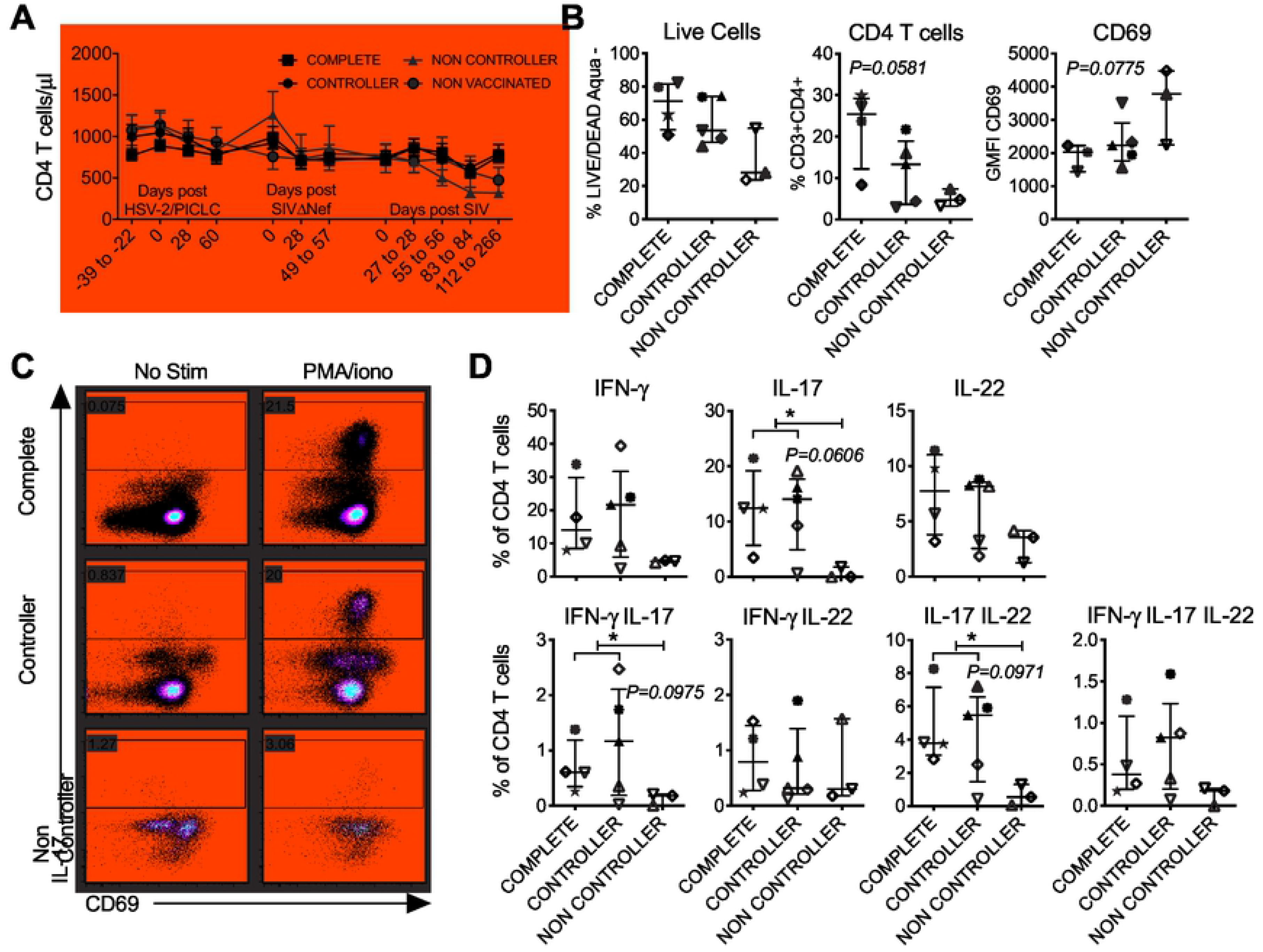
Control of SIV infection is associated with preservation of functional CD4 T cell subsets in the gastrointestinal tract. **(A)** Peripheral blood CD4 T cell counts are shown over time relative to treatment, vaccination, and SIV challenge for macaques at each level of protection. The mean ± SEM is shown for the animals in each protection group. **(B-D)** Functional profiling of T cells in gastrointestinal mucosa by flow cytometry in absence (No Stim) and presence (PMA/iono) of phorbol 12-myristate 13-acetate (PMA)/ionomycin stimulation. **(B)** Phenotype of unstimulated cells showing from left to right – the frequency of live cells (LIVE/DEAD AQUA^−^) within the singlet gate, frequency of CD4 T cells (AQUA^−^CD3^+^CD4^+^), and geometric mean fluorescence intensity (GMFI) of CD69 on CD4 T cells. Bars and whiskers indicate median ± IQR. **(C)** Flow cytometry for IL-17 and CD69 in unstimulated vs PMA/iono-stimulated cells in representative SIVΔNef-vaccinated macaques at each of the three levels of protection from SIV (complete protection, controller, and non-controller). **(D)** Frequency of cells secreting IFN-γ, IL-17, and IL-22 alone and in combination across the protection groups based on data from **(C)**. Bars and whiskers indicate median ± IQR. In **(B)** and **(D)**, differences between the 3 groups were assessed by Kruskal-Wallis test (*P=x* shown for *P<0.1* and considered significant for *P<0.05*). In **(D)**, comparison of non-controllers with the other groups was made using the Mann-Whitney test (* indicates Mann Whitney *P<0.05*). Throughout, treatment groups are color coded as in Figure 1 and symbols are as in Figure 1.

**Figure 4.**
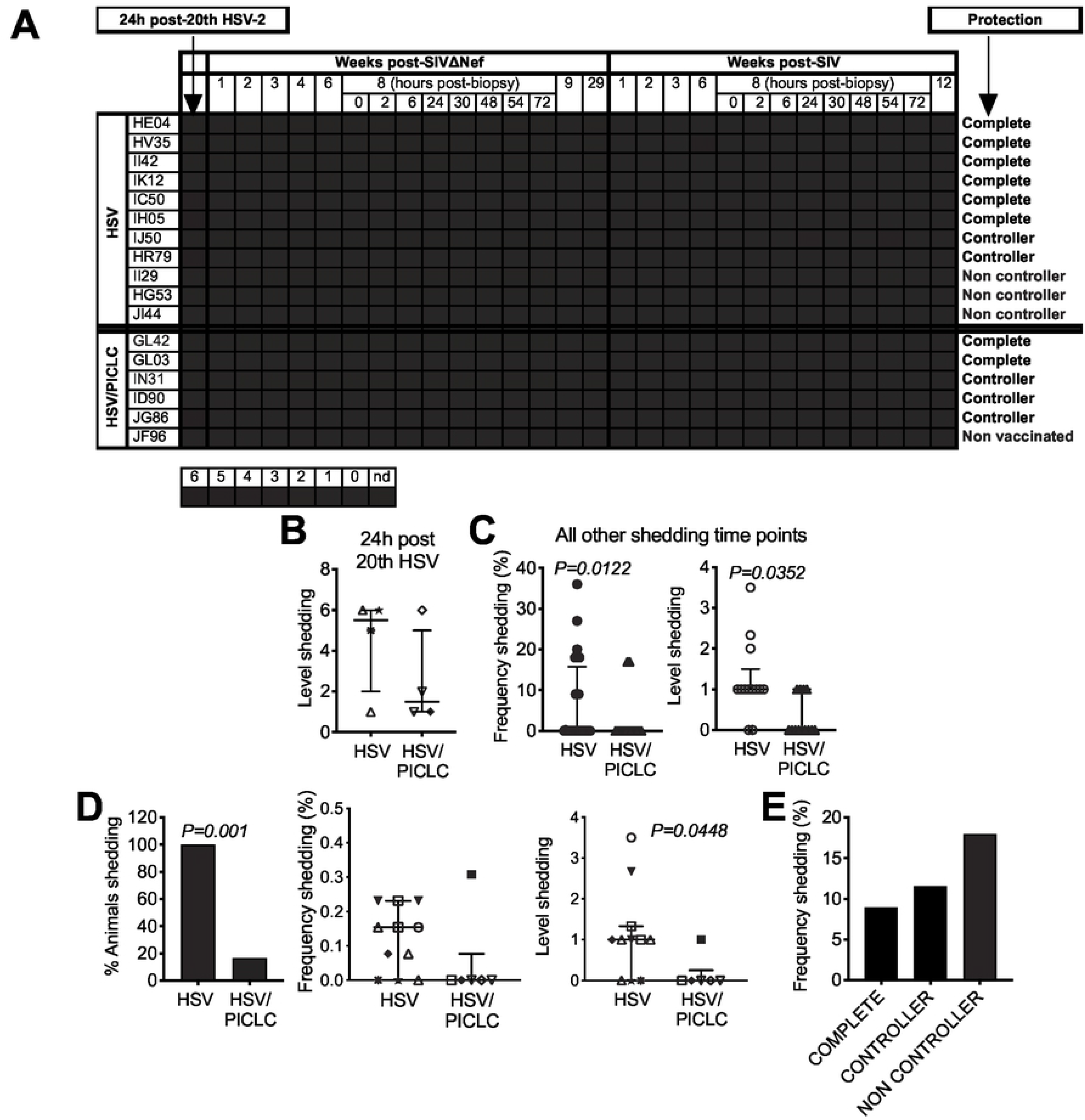
HSV-2 rectal shedding is decreased by polyICLC and tends to be increased with SIV pathogenesis. HSV-2 shedding in rectal mucosa was assessed in uncleared rectal swabs collected over the course of infection comprising both the LAV (Weeks post SIVΔNef) and SIV (Weeks post SIV) phases of the study. **(A)** Heat map depicting the relative quantities of HSV-2 gD detected in swabs over time by nested PCR (nPCR). Each row represents a macaque and each column a time point. The legend indicates the number of PCR reactions (of 6 total) that produced an HSV-2 amplicon. “nd” indicates that no sample was available for testing. **(B)** The relative level of shedding (number of amplicon-producing reactions of 6 total) detected in macaques 24 hours after the 20^th^ HSV-2/polyICLC exposure. Each symbol is a macaque; symbols are as in Figure 1. **(C)** Characteristics of shedding throughout the remainder of the study. Frequency of shedding (left) was calculated as the number of time points with shedding divided by all time points tested. Level of shedding (right) was calculated as the number of positive nPCR reactions divided by the number of time points on which shedding was detected. Each symbol represents a time point. **(D)** Shedding during the SIV phase: (Left) proportion of animals shedding at any post-SIV time point. (Center) Proportion of post-SIV time points with shedding. (Right) Number of PCR reactions with amplicon on post-SIV time points with shedding. Each symbol is an animal as in Figure 1. **(E)** Frequency of shedding during the SIV phase stratified by protection group. In **(B)**, **(C)**, and **(D)**, groups were compared by Mann Whitney test. In **(E)**, groups were compared by Chi squared test. P values <0.05 were considered significant.

**Figure 5.**
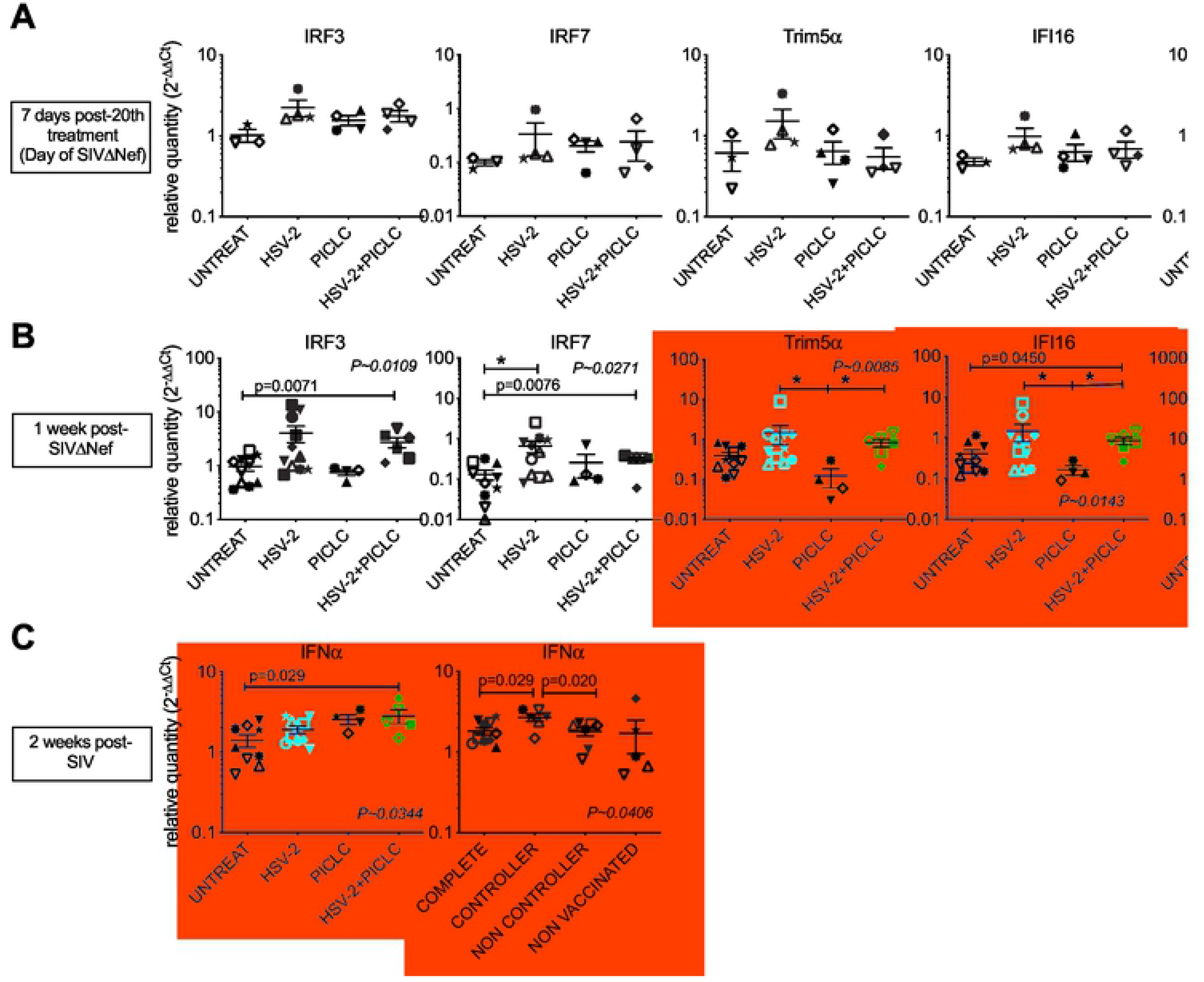
HSV-2 but not polyICLC elicits systemic type I IFN responses. Messenger RNA (mRNA) was measured in PBMCs for the genes indicated. Data are shown for **(A)** 7 days after the last treatment, which was the day of vaccination, **(B)** 1 week post-LAV, and **(C)** 2 weeks post-SIV. Bars and whiskers indicate mean ± SEM. Throughout, treatment groups are color coded as in Figure 1 and symbols are as in Figure 1. Comparison of all groups for each mRNA was performed using the Kruskal Wallis test (*P~x* shown for *P<0.05*) with Dunns post test for Kruskal Wallis *P<0.05*. * indicates Dunns P<0.05. The HSV-2/PICLC group was also compared with UNTREAT by Mann Whitney test. In **(C)**, controllers were compared with the other groups by Mann Whitney test.

**Figure 6.**
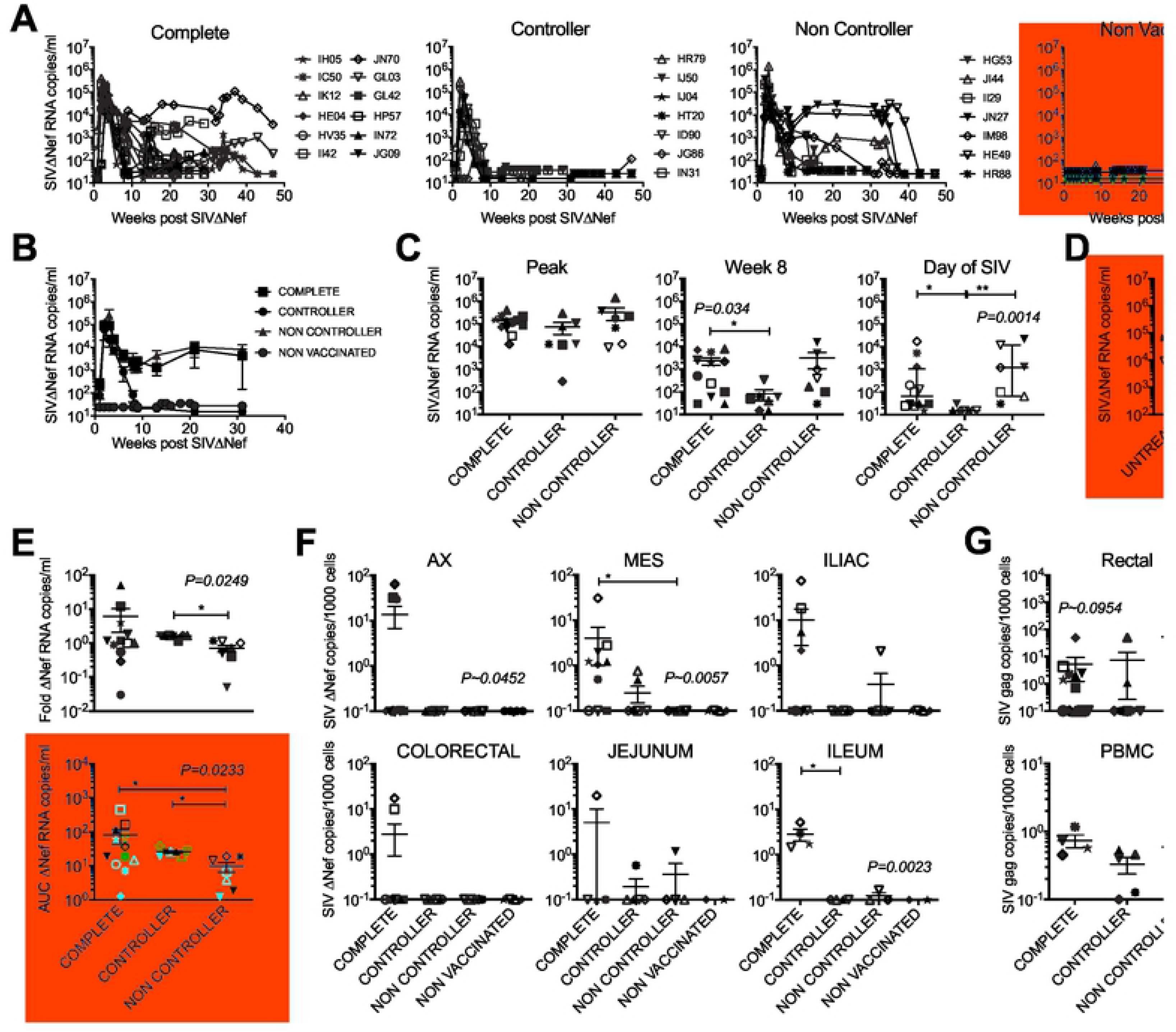
Absence of post-acute SIVΔNef viremia defines controller phenotype. **(A)** SIVΔNef loads in plasma of macaques according to their level of protection from SIV. Treatment groups are color coded as in Figure 1 and symbols are as in Figure 1. **(B)** Mean ± SEM for the SIVΔNef plasma levels of all animals at each level of protection. **(C)** Comparison of SIVΔNef viremia across protection groups at peak (highest viral load between weeks 2-4 post-vaccination, left), week 8 (center), and the day of SIV challenge (right). Each symbol is an animal as in (A). **(D)** Comparison of SIVΔNef plasma viral load across treatment groups at peak viremia. **(E)** SIVΔNef plasma viral load following SIV challenge in vaccinated macaques. Fold change within the first week after SIV challenge is shown on the top. Area under the curve (AUC) of the SIVΔNef plasma viral load during the 16 weeks of follow up post-SIV challenge is on the bottom. **(F)** Cell associated SIVΔNef DNA in tissues at necropsy. Tissues examined are the same as in Figure 1. **(G)** Cell associated SIVΔNef DNA in rectal tissue cells and PBMCs at 8 weeks post-LAV. In **(C-G)**, bars and whiskers indicate mean ± SEM. Comparisons between groups were made with Kruskal-Wallis test and Dunns post-test for Kruskal-Wallis P<0.05.

**Figure 7.**
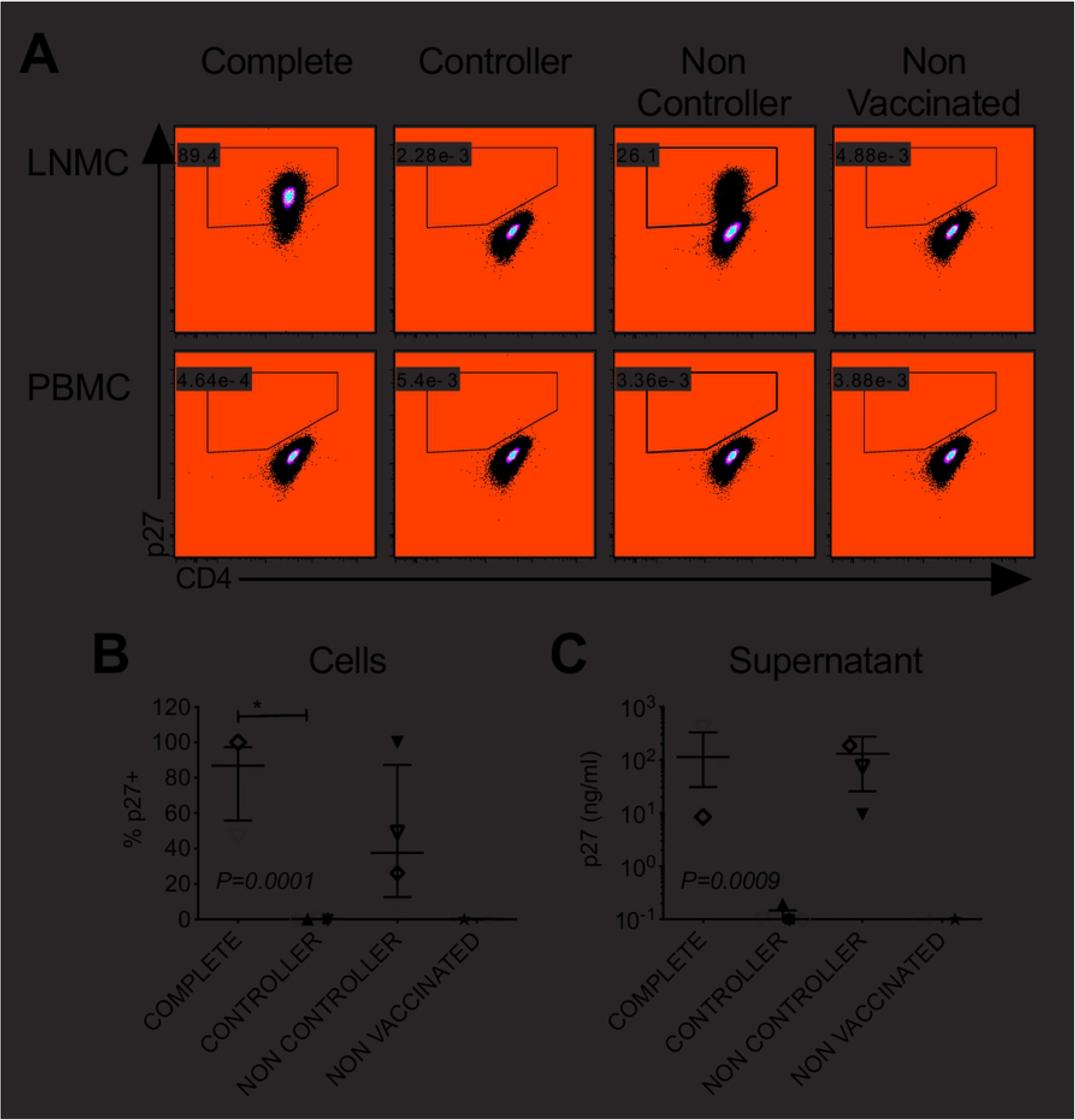
Persistent SIVΔNef replication in lymph nodes is not sufficient for protection. SIV growth was measured in co-cultures of CEMx174 cells with LNMCs or PBMCs from macaques 29 weeks post-LAV. Cells and supernatant were collected for analysis after 21 days of co-culture. **(A)** Flow cytometry plots from representative co-cultures from each protection group with axillary lymph node LNMCs (top) and PBMCs (bottom). Labeling for SIV gag p27 and CD4 is shown. **(B)** The percentage of p27+ cells is shown for all macaques tested (those challenged 31 weeks post-LAV). **(C)** p27 in the culture supernatant was quantified by ELISA. In **(B)** and **(C)**, bars and whiskers represent median ± IQR. Groups were compared by Kruskal Wallis test with Dunns post test for Kruskal-Wallis P<0.05.

**Figure 8.**
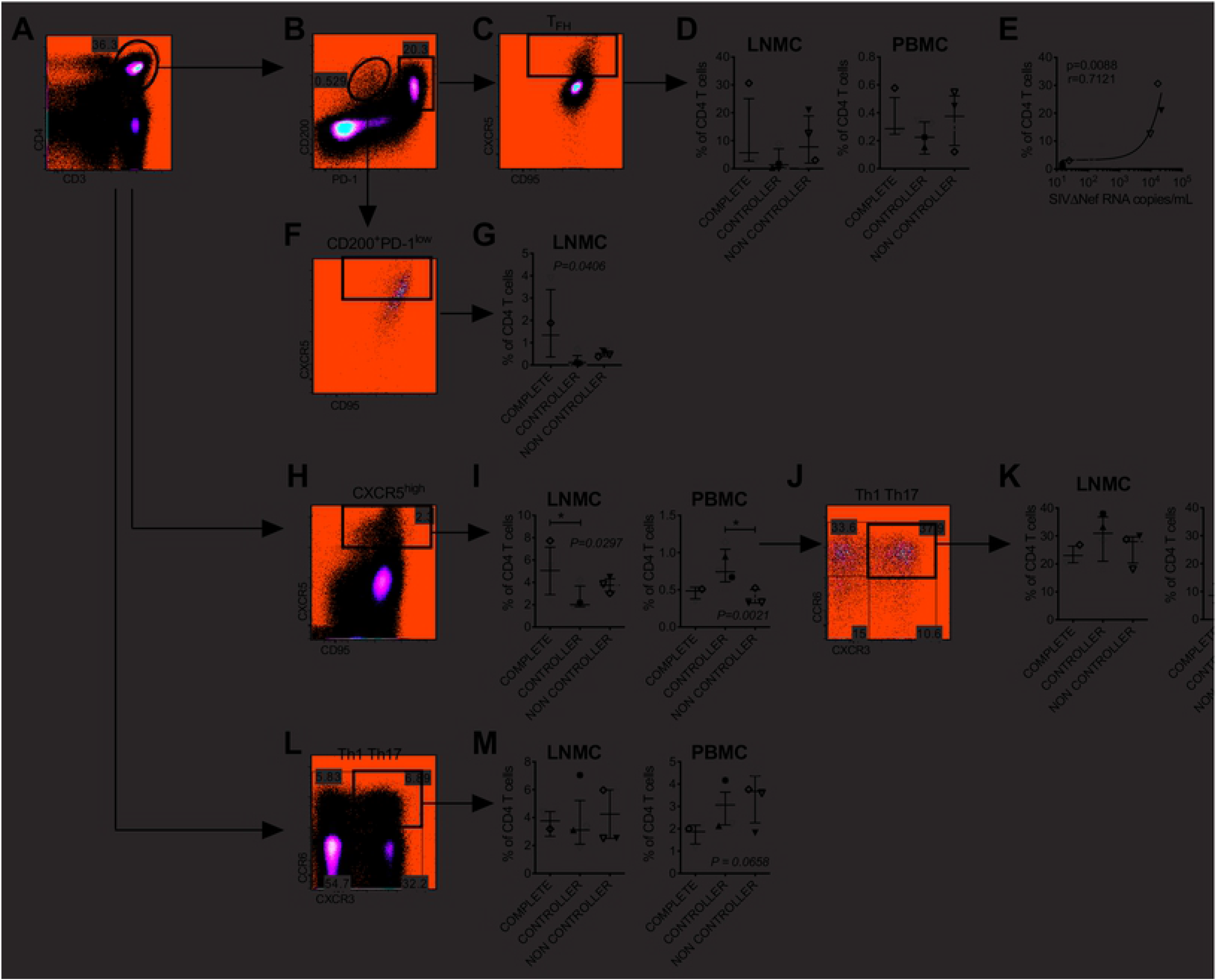
Pre-challenge CD4 T cell phenotype predicts protection outcomes. CD4 T cells within LNMCs and PBMCs from 29 weeks post-LAV were assessed by multicolor flow cytometry. **(A)** Gating of CD4 T cells by CD3 and CD4 expression within the live (LIVE/DEAD AQUA-) small cell singlet gate. **(B)** Gating of inguinal lymph node cells by PD-1 and CD200 expression. **(C)** CXCR5 and CD95 expression on PD-1^high^CD200^+^ Tfh in lymph node. **(D)** PD-1^high^CD200^+^ Tfh frequencies in LNMCs and PBMCs by protection group. **(E)** Correlation between Tfh frequency and concurrent SIVΔNef plasma viral load. Spearman r and p values are shown. **(F)** CXCR5 and CD95 expression on PD-1^low^CD200^+^ CD4 T cells from lymph node. **(G)** PD-1^low^CD200^+^ CD4 T cell frequencies in LNMCs by protection group. **(H)** CXCR5^high^ cells within total lymph node CD4 T cells. **(I)** CXCR5^high^ CD4 T cell frequencies in LNMCs and PBMCs by protection group. **(J)** Gating of CXCR5^high^ CD4 T cells from lymph node by CXCR3 and CCR6 expression. **(K)** Frequency of CXCR3^+^CCR6^+^ Th1Th17 cells within LNMCs and PBMCs by protection group. **(L)** Gating of CXCR3^+^CCR6^+^ cells within total lymph node CD4 T cells. **(M)** Frequency of CXCR3^+^CCR6^+^ CD4 T cells within LNMCs and PBMCs by protection group. Throughout, groups were compared by Kruskal Wallis test with Dunns post test unless otherwise noted. Bars and whiskers indicate median ± IQR. Treatment groups are color coded as in Figure 1 and symbols are as in Figure 1.

**Figure 9.**
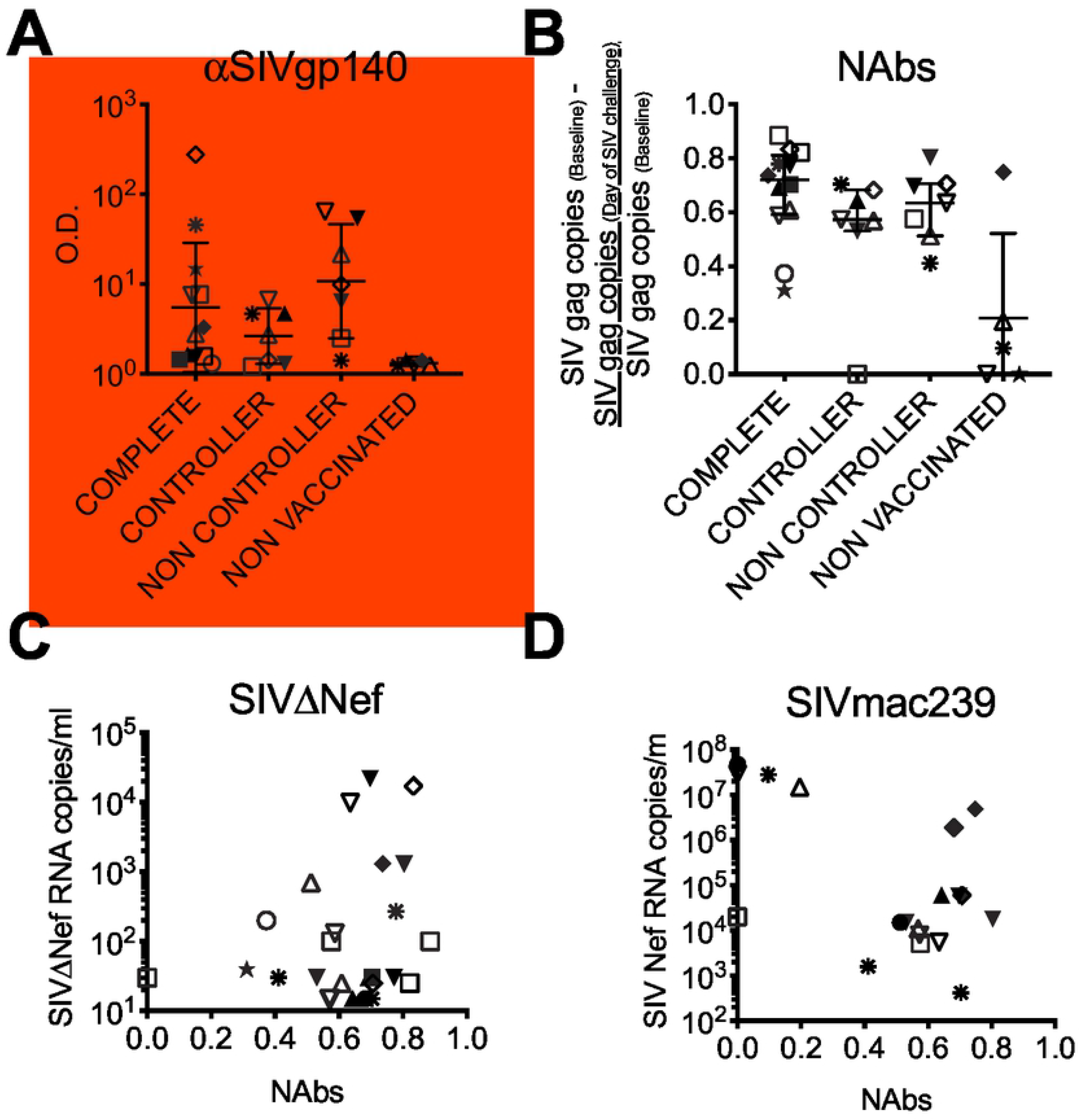
Neutralizing antibody frequency reflects the LAV load and does not predict protection. **(A)** The frequency of binding antibodies on the day of SIV challenge in plasma is shown for all 31 macaques in the study. Binding antibodies are quantified as the optical density (OD) in ELISA of plasma at a 1:2560 dilution binding to SIVmac1A11 gp140. **(B)** The frequency of neutralizing antibodies on the day of SIV challenge in plasma for 31 macaques. Neutralizing antibodies are quantified as the copy number of SIV *gag* detected in culture supernatants of CEMx174 cells in the presence of baseline plasma minus the copy number in the presence of immune plasma from day of SIV challenge divided by the copy number in the presence of baseline plasma. All plasmas were used at 1:640 dilution. **(C)** Correlation between neutralizing antibodies and the concurrent SIVΔNef plasma viremia. **(D)** Correlation between neutralizing antibodies and the peak SIV plasma viremia in animals that became SIV+.

In the untreated group, low-dose rectal immunization resulted in SIVΔNef infection (vaccine take, SIVΔNef+) in 60% of the macaques (**Table 1, Fig. 1B**), similar to our previous observations (15). Immunological stimulation of the rectal mucosa with HSV-2 or polyICLC prior to SIVΔNef exposure significantly enhanced vaccine take, increasing the proportion of SIVΔNef+ animals (**Table 1, Fig. 1B**). We then challenged all macaques rectally with 3×10^3^ TCID_50_ of virulent SIVmac239 and followed them for 16 weeks. Because both SIVΔNef and SIV are detected by standard quantification of gag RNA, we evaluated the effects of pre-vaccination treatment on protection by quantifying nef RNA to detect the intact Nef (in the challenge virus SIVmac239) or the region spanning the nef deletion (in the SIVΔNef LAV) (15, 17, 32). Overall, half (3/6) of the control group SIVΔNef+ macaques showed no evidence of SIVmac239 challenge virus infection (“complete protection”, defined by no time point with SIVmac239 plasma virus RNA >30 copies/ml plasma) while the other half became infected with levels of SIVmac239 plasma viremia similar to unvaccinated controls (**Table 1, Fig. 1C,D,E**). Complete protection was neither improved nor impaired significantly by mucosal stimulation (**Table 1, Fig. 1C,D,E**).

**Table 1.** Protection from SIVmac239 in SIVΔNef vaccinated macaques.

| Group | ΔNEF+ | Protection from WT |  |  |  |  |  |  |  |  |  |  |  |
| --- | --- | --- | --- | --- | --- | --- | --- | --- | --- | --- | --- | --- | --- |
|  |  | Complete |  |  | Controller |  |  | Any Protection |  |  | Non Controller |  |  |
|  |  | Total | 13<sup>a</sup> | 31<sup>a</sup> | Total | 13 | 31 | Total | 13 | 31 | Total | 13 | 31 |
| Untreated | 6/10 (60%) | 3/6 (50%) | 3 | 0 | 0/6 (0%) | 0 | 0 | 3/6 (50%) | 3 | 0 | 3/6 (50%) | 1 | 2 |
| HSV-2 | 11/11 (100%) | 6/11 (55%) | 4 | 2 | 2/11 (18%) | 1 | 1 | 8/11 (73%) | 5 | 3 | 3/11 (27%) | 2 | 1 |
| PICLC | 4/4 (100%) | 1/4 (25%) | 0 | 1 | 2/4 (50%) | 0 | 2 | 3/4 (75%) | 0 | 3 | 1/4 (25%) | 0 | 1 |
| HSV-2/PICLC | 5/6 (83%) | 2/5 (40%) | 1 | 1 | 3/5 (60%) | 1 | 2 | 5/5 (100%) | 2 | 3 | 0/5 (0%) | 0 | 0 |
| MHC Censored |  |  |  |  |  |  |  |  |  |  |  |  |  |
| Untreated | 5/10 (50%) | 2/5 (40%) | 2 | 0 | 0/5 (0%) | 0 | 0 | 2/5 (40%) | 2 | 0 | 3/5 (60%) | 1 | 2 |
| HSV-2 | 9/9 (100%) | 5/9 (55%) | 3 | 2 | 1/9 (11%) | 0 | 1 | 6/9 (67%) | 3 | 3 | 3/9 (33%) | 2 | 1 |
| PICLC | 1/1 (100%) | 1/1 (100%) | 0 | 1 | 0/1 (0%) | 0 | 0 | 1/1 (100%) | 0 | 1 | 0/1 (0%) | 0 | 0 |
| HSV-2/PICLC | 3/6 (50%) | 1/3 (33%) | 1 | 0 | 2/3 (67%) | 1 | 1 | 3/3 (100%) | 2 | 1 | 0/3 (0%) | 0 | 0 |

<sup>a</sup>Macaques in the column labeled "13" were challenged with SIV 13 weeks after SIVΔNef vaccination. Macaques in the column labeled "31" were challenged with SIV 31 weeks after SIVΔNef vaccination.

In contrast, mucosal stimulation by HSV-2, polyICLC, or both prior to vaccination was associated with the appearance of a partial protection “controller” phenotype that was absent in untreated animals (**Table 1, Fig. 1C,D,E**). Partial protection with relative control of viral replication (“controller”) was defined as SIVmac239 challenge virus infection with rapid post-acute containment and maintenance of plasma viremia <10^4^ RNA copies/ml (**Fig. 1D,E,F**). Two macaques that marginally fit these criteria were considered controllers – JG86 had higher peak viral load than the rest but experienced rapid decline in viremia and contained peripheral SIV replication to < 30 copies/mL by 16 weeks post infection (the last time point sampled); IJ50 had low peak viral load and rapid containment to less than 100 RNA copies/ml from weeks 4-8 post infection but then experienced resurgence of SIV plasma viremia at weeks 12 and 16 (**Fig. 1D**). Nonetheless, plasma SIV viral load in IJ50 was still a Log lower than in most non-controllers and several Logs lower than in unvaccinated macaques at week 16 (**Fig. 1F**); thus, IJ50 was grouped with the controllers. Controller status associated with protective MHC alleles was only seen in 25-50% of unvaccinated animals. The combination of HSV-2 and polyICLC resulted in the greatest frequency of controller macaques (**Table 1, Fig.1C,D**) and “any protection”, categorized as complete protection or control (**Table 1, Fig. 1C**). Although the controller phenotype was potentially due in part to MHC haplotype, when macaques with known protective Mamu alleles were censored, the controller phenotype was still most prevalent among HSV-2/polyICLC-treated macaques (**Table 1, Fig. S1, Fig. S2**), indicating a role for the stimuli. The impact of polyICLC alone on the controller phenotype could not be evaluated in the MHC censored dataset as only one polyICLC-treated macaque was negative for all three alleles. However, the controller phenotype was more prevalent within the HSV-2/polyICLC group than the HSV-2 group, suggesting a role for polyICLC (**Fig. S2**).

The study was executed in two parts (**Table 1, Fig. S1**). The first set of macaques was challenged 13 weeks post-SIVΔNef to mirror our previous study (15). Upon noting the emergence of the controller phenotype, we held the second set longer before challenge (31 weeks) to determine if this phenotype was related to incomplete maturation of the immune response 13 weeks post-SIVΔNef (10, 33, 34). However, in this low dose rectal SIVΔNef inoculation model, increasing the time between vaccination and challenge from 13 to 31 weeks neither increased the proportion of completely protected or controller macaques in the untreated group nor changed the course of viremia in SIVmac239 challenge virus infected macaques (**Table 1, Fig. S1**). This agrees with the shorter time to maturation of immunity against the homologous SIVmac239 vs heterologous challenge strains (35). Thus, we grouped animals from 13 and 31 weeks for further analyses.

Protection groupings based on plasma viral load were corroborated through measurements of SIVmac239 challenge virus DNA in lymphoid and mucosal tissues at the time of euthanasia, 16 weeks post-SIV challenge, assessed by quantitation of SIV Nef DNA (**Fig. 2**). As expected, completely protected macaques had no detectable SIVmac239 Nef DNA in axillary, mesenteric, or iliac lymph nodes, colorectal tissue, or gut mucosa. Non-controllers and unvaccinated macaques alike had high levels of SIV DNA in all tissues. Controllers varied in their level of tissue SIVmac239 Nef DNA with tissue SIV DNA viral load generally following plasma SIV RNA viral load (**Fig. 2, Fig. 1D,F**). IJ50 in the HSV-2 group was viremic with SIVmac239 Nef DNA detected at the time of necropsy, but SIVmac239 Nef DNA was only detected in the mesenteric lymph node among the tissues studied. In the MHC-censored dataset, the trends were the same but only one of the controllers had SIVmac239 Nef DNA detected in tissues at necropsy, the HSV-2/polyICLC-treated macaque ID90 (**Fig. S3**).

We further explored the degree of protection by examining CD4 T cell loss. Like completely protected macaques, controllers preserved their peripheral CD4 T cells while non-controllers lost over half of them, even more than non-vaccinated animals (**Fig. 3A**). In jejunum obtained from the SIV infected animals challenged 31 weeks post-SIVΔNef, we found that fewer live cells, fewer CD4 T cells among the live cells, and a more activated CD4 T cell phenotype (CD69+) tended to be present in animals with a lower degree of protection (**Fig. 3B**). Examination of the frequency of jejunum CD4 T cells secreting IFN-γ, IL-17, and IL-22 in response to mitogen stimulation revealed that controllers, but not non-controllers, also preserved their functional CD4 T cells, especially the Th17, Th1Th17, and Th17Th22 subsets in the gut at similar levels to the completely protected macaques (**Fig. 3C,D**). There was no difference in the frequency of CD8 T cells secreting the same cytokines (not shown). Peripheral T cell loss and gut T cell phenotype followed the same pattern in the MHC-censored dataset. Notably, the Th1 preservation was driven by animals with protective MHC alleles while IL-17 and IL-22 secreting cells were preserved in controllers without protective MHC alleles (**Fig. S4**). Overall, gut T cell function was somewhat reduced in the MHC-censored controllers compared to the total, suggesting a more prominent role for MHC-mediated protection in preserving gut T cell function. Taken together, these analyses indicate that both repeated low-dose rectal HSV-2 inoculation and repeated polyICLC treatment preceding SIVΔNef boosted vaccine uptake, and the combination of these stimuli in particular increased control of SIV replication in animals with breakthrough SIVmac239 challenge virus infection, which was associated with protection from gut CD4 T cell loss.

### Minimally virulent rectal HSV-2 infection elicits systemic type I IFN (IFN I) signaling that contributes to SIV control

Although rectal HSV-2 infection increased SIVΔNef infection (vaccine take), it did not reduce protection from wild type SIV and possibly improved protection, especially when administered together with polyICLC. Thus, we investigated HSV-2 shedding and innate immune responses in HSV-2 exposed macaques to appreciate the level of HSV-2 infection and understand if innate responses to HSV-2 could have aided protection from SIV. We detected HSV-2 DNA in rectal swabs from all but one of the HSV-2 exposed macaques (IN31 in the HSV-2/polyICLC group) on at least one of the time points sampled (despite lacking samples during the vaccination phase of those animals challenged 13 weeks post-SIVΔNef) (**Fig. 4A**). In macaques infected rectally with HSV-2 in the absence of polyICLC, shedding recurred infrequently during the study but over at least 25 weeks after the final HSV-2 inoculation, including in response to rectal mucosa biopsy especially in SIVmac239 infected animals (**Fig. 4A**).

PolyICLC significantly inhibited HSV-2 shedding over the course of the study. Twenty-four hours after the 20^th^ HSV-2 challenge, the level of HSV-2 DNA detected in rectal swabs was not significantly different between the HSV-2 and HSV-2/polyICLC groups, though it tended to be less in macaques treated with polyICLC (**Fig. 4A,B**). Analysis of all later time points revealed that animals treated with polyICLC shed HSV-2 significantly less frequently and had less HSV-2 DNA in their swabs at shedding times than those not treated with polyICLC (**Fig. 4A,C**). In fact, only one of the HSV-2/polyICLC animals (GL42) was shedding at any of the times examined past 24 hours (**Fig. 4A**). During just the post-SIVmac239 challenge phase (when samples were available from all animals) significantly fewer of the polyICLC treated animals experienced shedding, and the frequency and level of shedding in polyICLC treated animals were both reduced (**Fig. 4D**). In agreement with the greatest control of breakthrough infection with SIVmac239 seen in HSV-2/polyICLC treatment, shedding appeared to associate inversely with SIV protection status, especially in the non-polyICLC treated animals (**Fig. 4A**). During the period following SIV challenge, the frequency of shedding trended with the severity of SIV infection though too few macaques were studied to detect a significant effect **(Fig. 4E)**.

Further evidence that repeated low dose rectal HSV-2 inoculation resulted in a less virulent infection than high dose inoculation came by examining the impact of HSV-2 on the rectal mucosa. The macaques exhibited little acute rectal inflammatory response to repeated low dose HSV-2 (**Fig. S5**), which contrasts the response to a single high dose inoculation (17). In particular, CXCL8 was not elevated in swabs taken 24 hours after the final inoculation. Low dose rectal HSV-2 infection also failed to increase rectal T cell activation in biopsies collected 8 weeks post-SIVΔNef. Expression of CD69, CCR5, CCR6, CCR7, and α4β7 integrin on memory CD4 T cells were unaltered or potentially decreased by HSV-2 at this time point (**Fig. S5**), in contrast to what has been shown following single high dose mucosal HSV-2 inoculation (17, 24).

We next examined if HSV-2 infection had systemic effects despite minimal mucosal responses. In plasma 24 hours after the final treatment, levels of multiple inflammatory soluble factors were unaffected by HSV-2 though IL-2 was decreased by all treatments and CXCL9 and CCL11 were increased in plasma in association with the controller phenotype (ie. Highest level in HSV-2/polyICLC treated macaques) (**Fig. S5**). In PBMCs, we also studied gene expression relevant to the IFN I pathway, which is involved in the immune response to both HSV-2 and polyICLC. We quantified transcripts of two transcription factors (*IRF3* and *IRF7*) (36–38) involved in the induction of IFN responses to HSV infection and polyICLC, two IFN-induced innate antiviral proteins (*TRIM5α* and *IFI16*) associated with restriction of both HIV (39, 40) and HSV (41–43), and *IFNα*. Seven days after the last mucosal treatment (the day of vaccination), IFN I related transcripts were elevated in PBMCs isolated from HSV-2 infected macaques, especially *IRF3*, *TRIM5α*, and *IFI16*, whereas polyICLC had little effect (**Fig. 5A**). One week later when responses reflected both treatment and acute innate response to SIVΔNef, increased systemic expression of IFN I related genes was still related predominantly to HSV-2 infection, and polyICLC may have even dampened *TRIM5α* expression (**Fig. 5B**). However, *IRF3*, *TRIM5α*, and *IFI16* were all increased in the HSV-2/polyICLC group compared to Untreated macaques. Although IFN I responses at these time points did not correlate with eventual protection from SIV, at 2 weeks post SIV challenge, the expression of *IFNα* was increased in the controller group compared with either the completely protected or non-controller groups in parallel with the increased frequency of controller phenotype by treatment group (HSV-2/polyICLC > polyICLC > HSV-2 > Untreated) (**Fig. 5C**). Thus, polyICLC reduced HSV-2 shedding in the absence of a detectable systemic IFN I response at the times examined, whereas low-virulence HSV-2 infection itself triggered a systemic IFN I response despite little shedding or mucosal immune dysfunction at the times examined. Acute systemic IFNα production, which was elevated in macaques treated with mucosal stimuli, was associated with SIVmac239 control in SIV+ animals.

### SIVΔNef persists in lymph nodes of non-controllers but not controllers

In past studies using intravenous immunization with either intravenous or vaginal challenge, the measured levels of LAV, especially in lymph nodes, correlated with protection from virulent SIV challenge (6, 8, 44). We also previously found in our rectal immunization model that polyICLC increased levels of SIVΔNef viremia in association with increased protection from SIV (15). If SIVΔNef persistence was boosted through mucosal stimulation, this would provide a strong predictor of protection differentiating controllers from non-controllers and (together with increased acute IFNα) explain the controller phenotype.

In this unique model of SIVΔNef immunization across the rectal mucosa, we found that some of the completely protected macaques exhibited a transient SIVΔNef viremia throughout the time post-vaccination before SIVmac239 challenge (the vaccination phase) while others exhibited ongoing SIVΔNef viremia throughout the vaccination phase (**Fig. 6A,B**). In contrast, all controllers rapidly controlled SIVΔNef to <30 copies/mL plasma after a peak of replication in plasma (**Fig. 6A,B**). Non-controllers exhibited overlapping SIVΔNef plasma viral loads to the completely protected animals (**Fig. 6A,B**). Peak SIVΔNef viral loads were the same among the protected, controller, and non-controller groups, diverging in the post-acute period by 8 weeks post-vaccination through to the time of SIV challenge (**Fig. 6C**). Instead, peak SIVΔNef loads stratified by treatment; HSV-2 alone and not polyICLC (or the HSV-2/polyICLC combination) significantly increased SIVΔNef peak viral load (**Fig. 6D**). These trends were preserved in the MHC-censored dataset (**Fig. S6**).

Following SIV challenge, some of the completely protected macaques (those that had low to undetectable SIVΔNef viral load prior to SIV challenge) experienced an early resurgence of SIVΔNef RNA in plasma (**Fig. 6E**). By contrast controllers, which also had undetectable SIVΔNef viral load at the time of challenge, maintained SIVΔNef viremia <30 copies/mL plasma throughout the SIVmac239 challenge phase. Non-controllers exhibited an early and sustained decline in SIVΔNef regardless of their pre-challenge SIVΔNef viral load and concomitant with the development of SIVmac239 viremia. Paralleling their loss of SIVΔNef RNA from blood post-SIV challenge, non-controllers’ tissues contained no SIVΔNef DNA at the time of euthanasia 16 weeks post-challenge (**Fig. 6F**). Controllers also had little SIVΔNef DNA in tissues at euthanasia whereas completely protected animals had detectable SIVΔNef DNA in lymph nodes and mucosa, most prominently in the gut-draining mesenteric lymph nodes. Presence of SIVΔNef DNA in completely protected macaques was as expected based on prior association between SIVΔNef persistence and protection (8).

Lack of SIVΔNef DNA in tissues during chronic SIV infection in SIV+ macaques could have reflected replacement by SIV; alternatively, the loss of SIVΔNef in tissues could have preceded and facilitated SIV infection in non-controllers. To test the hypothesis that the extent of SIV protection was correlated with SIVΔNef persistence in tissues prior to SIV challenge, we examined ΔNef DNA in rectal biopsies taken 8 weeks post-vaccine. Unexpectedly, we detected similar copy numbers of SIVΔNef DNA between non-controllers and completely protected macaques (and more non-controllers than protected macaques had detectable SIVΔNef DNA, **Fig. 6G**). We also detected less SIVΔNef DNA in controllers, paralleling RNA levels in plasma.

In PBMCs from the same 8 weeks post-vaccine time point, SIVΔNef DNA levels followed the same pattern (**Fig. 6G**). These trends were also preserved in the MHC-censored dataset (**Fig S6**).

Since the presence of SIVΔNef DNA may not reflect ongoing SIVΔNef replication, we measured inducible replication competent virus in lymph nodes collected 29 weeks post-vaccination (2 weeks before SIV challenge) by culturing isolated lymph node mononuclear cells (LNMCs) with the susceptible CEMx174 cell line (**Fig. 7A**). We examined gag p27 production within cells (**Fig. 7B**) and secretion into culture supernatant (**Fig. 7C**). In accordance with the levels of pre-challenge SIVΔNef RNA measured in plasma and DNA in PBMCs and rectal mucosa, SIVΔNef grew in cultures containing LNMCs from both completely protected and non-controlling macaques. Of note, the growth tended to be lower overall in non-controllers than completely protected animals, especially when animals possessing protective MHC alleles were censored (**Fig. S7**), and the data represent a snapshot after 21 days of culture. Still it is noteworthy that no SIVΔNef grew in co-cultures containing LNMCs from controller macaques (**Fig. 7A,B,C**). No replication competent virus was detected in co-cultures containing PBMCs (**Fig. 7A, not shown**). These findings demonstrate that in this rectal model, SIVΔNef persists in blood plasma, PBMCs, mucosa, and lymph nodes of protected macaques. However, this persistence, which includes replication competent SIVΔNef in lymph nodes close to the time of SIV challenge, is not sufficient for complete protection from rectal SIV challenge. Neither is it necessary or sufficient for protection from disease.

### Lack of control predicted by pro-inflammatory, poorly functional T cell environment

We assessed the phenotype of CD4 T cells (**Fig. 8A**) in PBMCs and LNMCs from 2 weeks before SIV challenge with the goal to identify immunophenotypic patterns that distinguished completely protected macaques from non-controllers and to uncover novel phenotypes associated with control. Tfh cells (PD-1^high^CD200^+^ or PD-1^high^CXCR5^+^ CD4 T cells) within LNMCs have been shown to be a haven for HIV/SIV replication including SIVΔNef and SIV in elite controller macaques (8, 16). In our cohort, the frequency of cells with a Tfh phenotype in blood and lymph nodes (identified as PD-1^high^CD200^+^CXCR5^+^, **Fig. 8B,C**) correlated with SIVΔNef plasma viral load as expected (**Fig. 8D,E**). A population of PD-1^low^CD200^+^CXCR5^+^ cells having high CXCR5 expression was also identified in lymph nodes (**Fig. 8B,F**). These PD-1^low^ follicle homing cells, which have been shown to be Tfh precursors that support neutralizing antibody development (45), were present at highest frequency in the completely protected macaques (**Fig. 8G**). When we looked at all follicle-homing memory CD4 T cells (CD95^+^CXCR5^+^) with high CXCR5 expression (**Fig. 8H**), we found that these cells were more frequent in the blood of controllers than the other groups even though controllers had fewer Tfh cells, fewer PD-1^low^ Tfh precursors, and fewer CXCR5^high^ CD4 T cells in lymph nodes (**Fig. 8I**). These CXCR5^high^ cells in blood (but not lymph nodes) displayed an increasingly CXCR3^+^CCR6^+^ Th1Th17-like phenotype in macaques with less protection (**Fig. 8J,K**). There was no apparent difference in the frequency of either CXCR3^+^CCR6^−^ or CXCR3^−^CCR6^+^ cells between the groups (not shown). Similarly within the whole CD4 T cell population, macaques with less protection tended to have a greater frequency of CXCR3^+^CCR6^+^ cells in blood (**Fig. 8L,M**). These observations were similar in the MHC-censored dataset (**Fig. S8**).

Altered CXCR5^+^ CD4 T cell frequencies suggested a potential role for antibody maturation in the controller phenotype. However, when we measured the titer of SIV-specific antibodies in plasma on the day of challenge, we found that antibody titer followed SIVΔNef viremia (**Fig. 9A-C**) and did not correlate with peak SIVmac239 viremia in SIV+ macaques (**Fig. 9D, Fig. S9**).

We hypothesized that lack of control, which was associated with elevated Th1Th17 cell frequency, may be due to a dysregulated or inferior T cell response to SIV even though SIVΔNef was replicating in the non-controller macaques at the time of SIV challenge. We examined SIV-specific T cells 7 days post-SIV challenge, in order to observe boosted pre-existing responses without yet seeing *de novo* responses. Although SIV-specific T cell responses to SIV gag and env peptide pools were low, especially in the CD4 compartment (**Fig. S10**), we found in both the CD8 and CD4 T cell compartments that completely protected and non-controller macaques possessed a similar frequency of IFNγ-producing cells while controllers tended to have fewer of these cells, paralleling SIVΔNef persistence (**Fig. 10A,B, Fig. S10**). However, non-controllers’ T cells produced less TNFα than T cells from protected macaques, and IFNγ contributed most to their overall gag/env-specific CD8 T cell cytokine production (**Fig. 10C**). Similarly in the MHC censored data set, non-controllers had an IFNγ-dominant gag/env T cell response. In the full dataset, we noted that CD40L expression was minimal on SIV-specific T cells from non-controllers compared with controllers and completely protected macaques, most notably in the CD8 compartment (**Fig. 10B**). However, CD40L effects in controllers were related to protective MHC alleles, and when the animals possessing these alleles were censored, controllers and non-controllers both exhibited low CD40L on their antigen-specific T cells in comparison with completely protected animals (**Fig. S11**). Thus CD40L on CD8 T cells may contribute to the difference between controllers and completely protected macaques. Low CD40L expression in SIV+ animals was not global as PMA/ionomycin-stimulation increased CD40L in all animals similarly (**Fig. 10D**).

**Figure 10.**
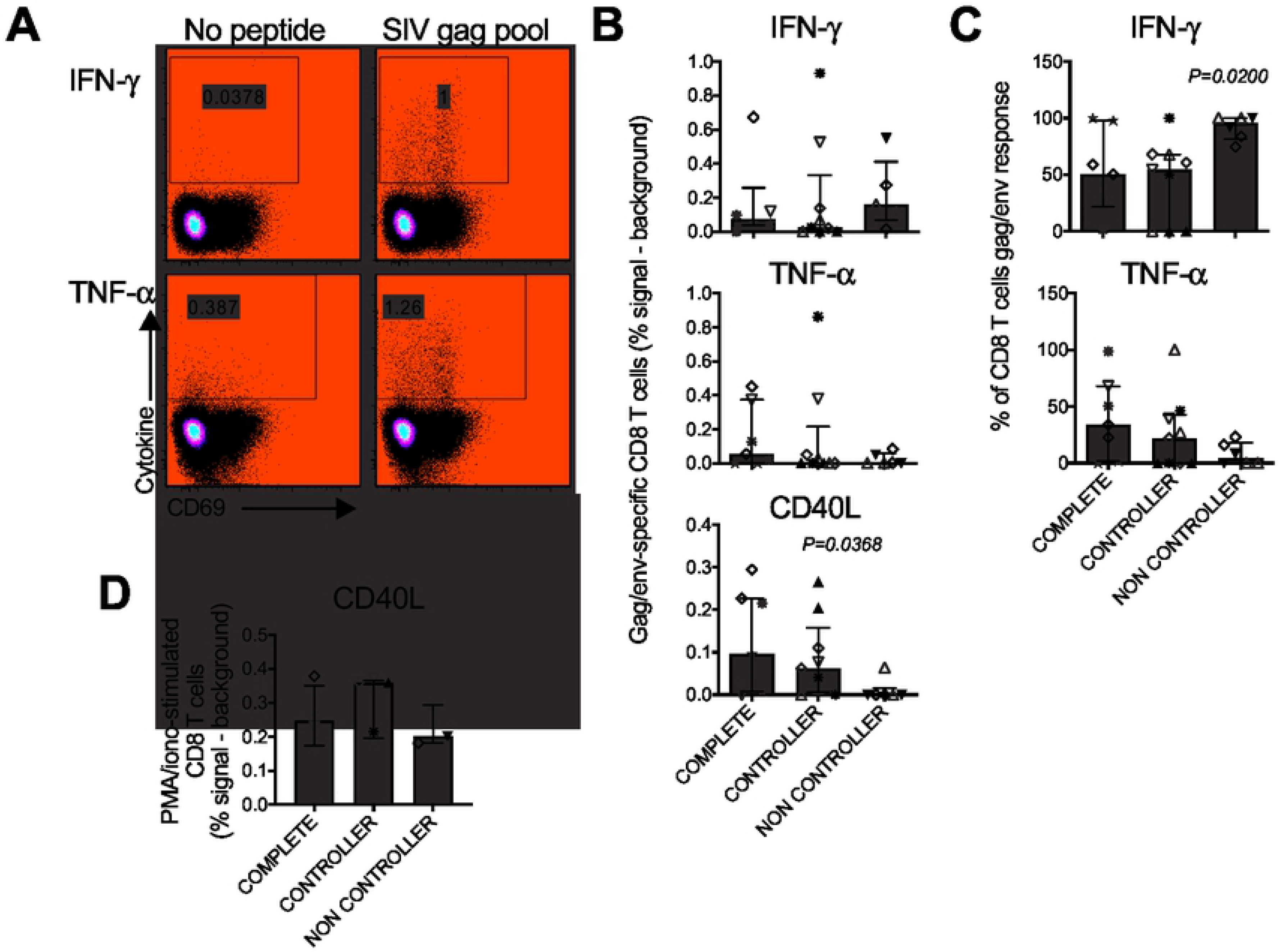
Polyfunctionality and a CD8 helper phenotype correlate with protection outcomes. The frequency and phenotype of SIV gag- and env-specific T cells were analyzed by intracellular cytokine flow cytometry using LNMCs isolated 7 days post-SIV challenge from axillary lymph nodes of macaques challenged 31 weeks post-LAV. Data for CD8 T cells are shown. **(A)** Production of IFNγ (top) and TNFα (bottom) in response to SIV gag peptide pool vs media (No peptide) shown for a representative macaque. Production of IL2 is not shown. **(B)** Frequency of CD8 T cells expressing the indicated cytokine or surface marker in response to gag pool and env pool in each macaque shown by protection group. Each macaque is shown twice – once for gag response and once for env response. Colors and symbols are as in Figure 1. **(C)** Proportion of total CD8 T cell response attributable to each cytokine. Each macaque is shown twice, indicating the gag- and env-specific responses. **(D)** CD40L surface expression on CD8 T cells in response to PMA/ionomycin stimulation.

## DISCUSSION

Adjuvants are a key element of successful vaccines and can promote increased vaccine efficacy by inducing certain kinds of immune activation. Yet paradoxically, agents that induce mucosal immune activation may undermine the efficacy of HIV vaccines because immune activation can increase HIV susceptibility by generating and recruiting permissive target cells for the virus. We set out to evaluate the impact of opposing triggers of immune activation on the SIVΔNef vaccine using a mucosal vaccination/challenge model. We hypothesized that HSV-2 infection would impair SIVΔNef-mediated immune responses whereas polyICLC would boost immune responses and potentially limit postulated HSV-2 mediated dysfunction. Unexpectedly, we found that neither HSV-2 nor polyICLC impacted complete protection by SIVΔNef but both promoted partial control of breakthrough SIVmac239 infections in SIVΔNef vaccinated animals, with the most pronounced control in animals that received both immune modulating stimuli in conjunction with SIVΔNef vaccination.

HSV-2 infection inflames the mucosa with cytokine secretion and recruitment of immune cells and impairs the immunostimulatory capacity of DCs to induce T cell responses (46–52). The IFN I pathway is involved in initial control of HSV and SIV infections (53), and IFN-α was shown to blunt SIV infection (53) while plasma IFN-α during chronic infection is also a hallmark of HIV/SIV systemic immune activation (54–60). IFN I responses are also triggered by polyIC and polyICLC (15, 18, 19, 61, 62), and polyIC was shown to blunt HSV-2 infection in a mouse genital infection model (63, 64). As used in our study, low dose repeated mucosal HSV-2 exposure elicited a minimal mucosal innate response even though macaques shed the virus intermittently throughout the follow up period. Negligible mucosal inflammation may explain why HSV-2 infection did not undermine SIVΔNef mediated protection. The repeated inoculation regimen with 10^7^ pfu HSV-2 per challenge was based on our previous finding that this approach augmented shedding following vaginal infection in contrast to a single inoculation with 2 x 10^8^ pfu (29, 30). However, in that study, HSV-2 was inoculated as a co-challenge with SHIV vaginally, and that may have facilitated more frequent and greater magnitude HSV-2 shedding. While it may be that HSV-2 infection is less virulent in macaques rectally than vaginally, previous studies found that rectal single high dose HSV-2 inoculation resulted in detectable mucosal and systemic inflammation in contrast to our observations herein (17, 24). It is notable that HSV-2 infection increased SIVΔNef infection and peak viremia despite the low virulence and absence of rectal cytokine responses or T cell activation at the times examined, underscoring that factors beyond mucosal inflammation are involved in the HSV-2 mediated increase in HIV/SIV susceptibility and replication. However, we did not study any other time points or examine a role for additional mucosal immune changes (e.g. HSV-specific tissue resident memory T cells) since we wanted to limit the collection of tissues.

PolyICLC also contributed to SIV control in SIVΔNef vaccinated macaques, as the frequency of controllers was higher in the HSV-2/polyICLC group than in the HSV-2 group even with the protective MHC allele-bearing macaques censored. PolyICLC and its parent molecule polyIC stimulate innate immunity, activate DCs to focus Th1 responses, and are being tested in preventative and therapeutic vaccines for HIV, other pathogens, and cancer in animal models and clinical trials (15, 19, 20, 62, 65–74). Repeated administration of polyICLC increased SIVΔNef take but did not increase SIVΔNef viremia or induce a systemic IFN I response detectable at the times tested. PolyICLC did, however, induce mucosal and systemic innate antiviral cytokine responses. We previously reported that a regimen of 2mg of polyICLC given twice 24 hours apart (with the second dose 24 hours before SIVΔNef) dampened take of SIVΔNef, increased SIVΔNef viremia in those vaccinated, and boosted complete protection from rectal SIV challenge (15). Differences in SIV protection outcomes by the dosing and timing of the same adjuvant highlight the delicate balance required in the mucosa for prevention of HIV transmission. Facilitation of complete protection by polyICLC may be linked with the magnitude of SIVΔNef viremia as we did not observe increased SIVΔNef viremia by polyICLC herein. In keeping with published results from murine studies showing that polyICLC inhibits HSV-2 infection (63, 64), polyICLC reduced HSV-2 shedding in the macaques. Since delivering polyICLC with each HSV-2 inoculation resulted in virtually no shedding past the 24-hour time point, it is likely that polyICLC either blunted the inoculum or triggered an abortive infection with immune system activation. Mechanistic studies that can reveal how the combination of HSV-2 and polyICLC uniquely focused the immune system for SIV control will be critically informative.

Most studies of SIVΔNef and other LAVs have used traditional intravenous immunization with a large vaccine inoculum (e.g. 10^5^ TCID_50_ SIVΔNef). In those studies, vaccine take was universal, and protection correlated inversely with the level of vaccine attenuation and heterogeneity between the vaccine and challenge strains (6–8, 33, 34). Immunization with 10^3^ TCID_50_ SIVΔNef rectally is less protective (15), providing the opportunity to investigate the role of host factors and immune responses to the same vaccine at differing levels of protection. The large cohort of macaques immunized mucosally with SIVΔNef and displaying varying levels of protection herein enabled the unexpected finding that in this mucosal vaccination model, SIVΔNef persistence – as reflected by levels of plasma viremia, SIVΔNef DNA in tissues, and replication competent virus in lymph nodes – is neither necessary nor sufficient for protection from SIV disease. In fact, macaques with the greatest SIVΔNef viremia exhibited no control whatsoever over SIVmac239 replication while macaques that controlled SIV completely after infection had no SIVΔNef – including DNA, RNA, and replication competent virus – in blood or tissues prior to SIV challenge. Ours is the first study of which we are aware to identify a specific SIVΔNef vaccinated partial protection group with complete control of SIV viremia in the absence of measurable SIVΔNef persistence. Although low amounts of SIV DNA were detected in the gut and lymph nodes of controller macaques at the time of euthanasia, CD4 T cell frequency and function in the gut mucosa were preserved in keeping with previous findings that completely and partially protected SIVΔNef vaccinated macaques do not lose their gut CD4 T cells (6). We further demonstrated that functional cytokine producing cells were preserved, especially the IL17-secreting subsets.

How vaccinated macaques managed to control SIV in the absence of SIVΔNef persistence remains mechanistically unanswered. Local immune changes, including innate responses at times we did not sample and migration or phenotypic/functional changes in specific T cell subsets as mentioned above, could have participated in the controller phenotype. Rectal HSV-2 infection did induce a systemic IFN I response, which was sustained in the polyICLC-treated animals at the times examined, and this also potentially contributed to SIV control. We explored a role for Tfh cells since previous studies identified Tfh cells in lymph nodes as the major refuge for LAV replication in vaccinated macaques as well as for SIV persistence with viral transcription in elite controllers (8, 16). Tfh cell frequency herein paralleled SIVΔNef viremia, which is not surprising since these cells are expected to house replicating virus. Although controller macaques had a low frequency of Tfh cells (paralleling lack of SIVΔNef viremia), they had a heightened frequency of CXCR5^high^ CD4 T cells in blood, suggesting a possible redistribution of immune cells between blood and tissues. A lower baseline frequency of Tfh cells could have been the cause for lower SIVΔNef persistence in controllers. By the same reasoning, a higher frequency of total CXCR5^high^ CD4 T cells in blood could have resulted from lack of SIVΔNef replication and thus lack of cell death. However, IFN I has been shown to induce the CXCR5 ligand CXCL13 in the periphery, driving the formation of extra-lymphoid germinal centers (75). Controllers also displayed increased IFNα production during acute SIV infection. The enhanced IFN I production by controllers (especially in HSV-2/polyICLC treated animals) could have independently facilitated the induction of CXCR5 on blood CD4 T cells. Future studies will need to investigate further the role of these blood CXCR5^high^ cells in SIV control.

Tfh frequency does not explain the differences in protection outcomes between the protected and non-controller macaques. Similar Tfh cell frequencies pre-challenge are likely related to the similar SIVΔNef viral loads. Importantly, we were unable to dissect which cells harbored SIVΔNef pre-challenge as co-cultures of LNMCs with CEMx174 cells utilized unsorted LNMC populations and the frequency of SIVΔNef+ cells was below the level of detection in absence of co-culture. But it is possible that different cell subsets from completely protected and non-controller macaques harbored the SIVΔNef pre-challenge and that this contributed to the dysfunctional immune response in non-controllers. Completely protected animals and not non-controllers exhibited a blip in SIVΔNef viremia following SIV challenge, but we do not know which cells produced this blip or if it aided protective immunity.

Non-controllers segregated from completely protected macaques phenotypically by their heightened frequency in blood of CXCR3+CCR6+ CD4 T cells, a subset containing the pro-inflammatory and highly HIV-susceptible Th1Th17 cells, within total CD4 T cells and also within the CXCR5^high^ subset of CD4 T cells. The correlation between Th1Th17 cell frequency and lack of control hints at the importance of systemic immune activation in HIV pathogenesis. Non-Controllers also lacked the PD-1^low^CXCR5^high^ Tfh precursors found in the protected macaques.

Location, maturity and functionality, especially of SIV-specific T cell responses, have emerged as cornerstones of SIVΔNef immunity (6, 8, 9, 44, 76, 77), and T cell function certainly was involved in the difference between completely protected and non-controller macaques. Previous studies showed the importance of lymph node T cell functionality in LAV-mediated protection in terms of the magnitude and number of cytokines secreted. Adding to this, we found that non-controllers also displayed an impaired CD8 T cell response dominated by monofunctional IFNγ producing cells. Both non-controllers and controllers differed from the completely protected macaques by having low CD40L expression on their SIV-specific T cells, and thus low capacity to receive co-stimulatory signals. Although CD40L is prototypically considered a marker of antigen specific CD4 T cells, CD40L expression has been documented on a substantial fraction of CD8 T cells with helper cytokine secretion function (78). Our findings indicate that the ability to engage costimulatory CD40 on cognate antigen presenting cells is an additional important factor in SIVΔNef mediated protection that may help to differentiate completely protected macaques from those not completely protected (controllers and non-controllers). Although HIV-induced impairment in CD40L expression has been reported, it is unlikely that SIV replication drove the lack of CD40L expression so early in infection. Whether impairment in CD40L was a contributing cause of failure to be completely protected from SIV infection remains to be investigated.

There were limitations to our study in terms of specific immune responses and host characteristics that were beyond the scope and so not measured herein. We did not determine the tetherin, APOBEC3G, or other alleles that could have promoted certain protection phenotypes. We did not study gene expression over time, but only at a single time point. We did not study antibody functions other than neutralization. And we did not examine SIV-specific T cells in the mucosa or HSV-specific T cells at all. Any of these parameters could have additionally contributed to the phenotypes we identified and should be examined in future studies. In addition, we followed animals out to only 16 weeks post SIV challenge, and the protection groupings were made based on the viral load data during this time period. Importantly, we are unable to exclude potential confounding of protective Mamu alleles as macaques possessing these alleles were included in the study. In particular, the effect of polyICLC alone on the controller phenotype cannot be evaluated as three of four polyICLC treated macaques possessed a protective allele. Nonetheless, the findings for HSV-2 persisted in the MHC censored dataset and the controller phenotype was present more in the HSV-2/polyICLC group than the HSV-2 group, indicating a role for polyICLC, and also the combination of the two agents. Moreover, most of the immunological observations were consistent between the full dataset and the MHC censored dataset. A central caveat to this work is that, despite enhancing SIVΔNef acquisition and initial replication, the HSV-2 infection resulting from low-dose repeated inoculation was minimally pathogenic. Hence, we could not explore how an underlying robust HSV-2 infection (as seen in humans) would influence protection by HIV vaccines. Use of the vaginal HSV-2 infection model, which leads to greater shedding even in the absence of SIV infection (22, 23, 29, 30), may be more useful and just as relevant for answering this question. A better understanding of how HSV-2 infection differs between macaques and humans may also facilitate an understanding of how to capitalize on the adjuvant properties of HSV observed in macaques and turn vaccination of HSV infected subjects to the favor of immunogenicity. Upon introduction of an HIV vaccine into the field, many adolescents and adults who undergo HIV immunization will already be infected with HSV-2, most with subclinical infection. Thus, understanding how underlying subclinical HSV-2 infection will influence protection is an important step in successful HIV vaccine development that should be pursued in future studies.

## MATERIALS AND METHODS

### Ethics statement

Adult male Indian rhesus macaques (*Macaca mulatta*) that tested negative by serology and virus-specific PCR for SIV, SRV, Herpes B, and STLV-1 were selected for these studies. Animal care at Tulane National Primate Research Center (TNPRC, Covington, LA) complied with the regulations stated in the Animal Welfare Act and the Guide for the Care and Use of Laboratory Animals (79, 80). All macaque studies were approved by the Institutional Animal Care and Use Committee (IACUC) of TNPRC for macaques (OLAW Assurance #A4499-01) and complied with TNPRC animal care procedures. TNPRC receives full accreditation by the Association for Accreditation of Laboratory Animal Care (AAALAC #000594). Animals were socially housed indoors in climate-controlled conditions and were monitored twice daily by a team of veterinarians and technicians to ensure their welfare. Any abnormalities, including changes in appetite, stool, and behavior, were recorded and reported to a veterinarian. Macaques were fed commercially prepared monkey chow twice daily. Supplemental foods were provided in the form of fruit, vegetables, and foraging treats as part of the TNPRC environmental enrichment program. Water was available continuously through an automated watering system.

Veterinarians at the TNPRC Division of Veterinary Medicine have established procedures to minimize pain and distress through several means in accordance with the recommendations of the Weatherall Report. Prior to all procedures, including blood draws, macaques were anesthetized with ketamine-HCl (10 mg/kg) or tiletamine/zolazepam (6 mg/kg). Preemptive and post-procedural analgesia (buprenorphine 0.01 mg/kg) was administered for procedures that could cause more than momentary pain or distress in humans undergoing the same procedures. Macaques were euthanized at the study conclusion 16 weeks post-SIVmac239 challenge using methods consistent with recommendations of the American Veterinary Medical Association (AVMA) Panel on Euthanasia and per the recommendations of the IACUC. Four macaques were recommended for euthanasia prior to study conclusion – 2 were euthanized prior to scheduled SIV challenge (and were excluded from the study analyses) and 2 were euthanized at 12 weeks post-SIV challenge. For euthanasia, animals were anesthetized with tiletamine/zolazepam (8 mg/kg) and given buprenorphine (0.01 mg/kg) followed by an overdose of pentobarbital sodium. Death was confirmed by auscultation of the heart and pupillary dilation.

### Viruses and stimuli

Stocks of SIVmac239 and SIVmac239ΔNef were grown in freshly isolated rhesus macaque PBMCs from SIV-uninfected single donors as previously described (15). The same virus stocks used in (15) were re-titered and used herein. Virus titer was determined in CEMx174 cells (ATCC, Manassass, VA) by p27 ELISA quantification (ZeptoMetrix, Buffalo, NY) and syncytia scoring after 14 days with the calculation method of Reed and Meunch. HSV-2 strain G was originally obtained from ATCC and grown and titered on Vero cells (ATCC) by plaque assay as described (81). PolyICLC (Hiltonol®) was provided by Oncovir (15, 19, 20, 65).

### Animal treatments and specimen collection

Thirty-three macaques were initially enrolled across four treatment groups: control (PBS; n=11), HSV-2 alone (n=11), polyICLC alone (n=5), and HSV-2/polyICLC (n=6). However, two macaques (one control, one polyICLC) became ill for reasons unrelated to the assigned treatments, had to be euthanized prior to scheduled SIV challenge, and were excluded from all analyses. Treatments were administered atraumatically rectally in 1 mL volume repeated twice weekly for 10 weeks (20 treatments) according to the protocol we developed for vaginal HSV-2 infection (29, 30). Each HSV-2 exposure was 10^7^ pfu; each polyICLC exposure was 1 mg; and each HSV-2/polyICLC exposure was 10^7^ pfu + 1 mg mixed together. Seven days after the last treatment, all macaques were rectally immunized with the LAV SIVΔNef (10^3^ TCID_50_). Half of the animals (IK12, HE04, HV35, II42, GL42, IN72, JG09, HP57, IJ50, IN31, HR88, HG53, II29, IR67, II95, HP47) were challenged with SIV 13 weeks later and the other half (IC50, IH05, GL03, JN70, HR79, ID90, JG86, HT20, IJ04, JI44, HE49, IM98, JN27, JM05, JF96) 31 weeks later (**Fig. S1**). Challenging macaques that did not become infected with SIVΔNef alongside vaccinated macaques provided an internal control for SIV virulence (all unvaccinated animals became infected with SIV). The animals were followed for 16 weeks post-SIV challenge and euthanized.

Blood, rectal swabs, rectal biopsies, and peripheral lymph nodes (inguinal, axillary) were collected periodically during the study. At euthanasia, additional deep tissues (axillary, mesenteric, iliac lymph nodes; colorectal mucosa; jejunum; ileum) were collected. All fluids and tissues were shipped to the Population Council in New York overnight and processed immediately on arrival as previously described (15, 19). Plasmas were isolated and stored at −80°C (15). Isolated PBMCs were used immediately for flow cytometry or stored in RNA Protect (Qiagen) according to the manufacturer’s instructions for RNA isolation. Rectal swabs were stored both as total uncleared and cleared (by centrifugation) aliquots at −80°C (15, 17). Mucosal tissues and lymph nodes were transported in L-15 media (HyClone Laboratories, Inc., Logan, UT) supplemented with 10% FBS and 100 U/mL penicillin/100 μg/mL streptomycin. All tissues were washed upon arrival. Lymph nodes were manually dissociated and the LNMCs passed through 40μm cell strainers before being used for flow cytometry, or they were placed in RNALater (Qiagen) overnight at 4°C before being transferred to −20°C for storage. Mucosal tissues were digested with collagenase IV and passed through 40μm strainers to obtain cells for flow cytometry, or they were placed in RNALater and stored.

### SIVΔNef and SIV detection

SIVΔNef and SIV plasma RNA viral loads in macaques were determined by discriminatory RT-qPCR in *nef* as previously described (15, 17, 32). The Quantitative Molecular Diagnostics Core, AIDS and Cancer Virus Program, Frederick National Laboratory performed the plasma Nef and ΔNef assays. Infection was defined as two consecutive time points with plasma viremia >100 copies/mL or any viremia >10^3^ copies/mL, consistent with our previously defined criteria (15). SIVΔNef and SIV DNA in tissues collected at the time of euthanasia were quantified alongside albumin by qPCR in-house (17, 82, 83) using the Nef and ΔNef-specific primers and probes. Standard curves for the DNA qPCR assays were produced from SIV and SIVΔNef virus preparations. RNA was extracted from SIV and SIVΔNef virus stocks with the QIAamp UltraSens Virus kit (Qiagen, Germantown, MD), reverse transcribed with the Superscript VILO kit (Thermofisher, Waltham, MA), and diluted to make the standard curves. Each standard curve was checked for specificity and amplification linearity across the dilutions.

### HSV-2 detection

We determined the presence of HSV-2 DNA in unclarified rectal swab samples collected over time, including following biopsy of the rectal mucosa 8 weeks post-SIVΔNef and 8 weeks post-SIV. The biopsy provides a stressor to encourage HSV-2 reactivation and mucosal replication, thereby increasing mucosal shedding. Swabs were subjected to nested PCR with 6 reactions per sample, as in previous studies (17, 22, 24, 29, 30, 82, 83). The identity of the amplicons was confirmed by sequencing (Genewiz, South Plainfield, NJ).

### Soluble factors

Soluble factors in rectal swabs and plasma were quantified using the monkey Novex Multiplex Luminex assay (Cytokine Monkey Magnetic 29-Plex Panel; Invitrogen, Waltham, MA) on a MAGPIX1 System (Luminex Corporation, Austin, TX). The kit included the following analytes: IL-1RA, GM-CSF, G-CSF, MDC, MIF, I-TAC, FGF-Basic, EGF, HGF, VEGF, Eotaxin, TNFα, IFNγ, IL-1β, IL-2, IL-4, IL-5, IL-6, IL-10, IL-12, IL-15, IL-17, CXCL8, CXCL9, CXCL10, CCL2, CCL3, CCL4 and CCL5. Factors with detectable levels above the lowest standard curve concentration were analyzed.

### Gene expression

Gene expression studies were performed with modifications to our previously published approach (15, 84). RNA was isolated from frozen PBMC dry pellets using the RNeasy mini kit (Qiagen) according to the manufacturer’s instructions with Qiashredder columns (Qiagen) for cell disruption. Total RNA was subjected to on-column DNA digestion with RNase-free DNase (Qiagen) and post-isolation DNA digestion using DNA-free DNase Treatment and Removal System from Ambion (Austin, TX) according to the manufacturer’s instructions. RNA was quantified on a Nanodrop 1000 spectrophotometer (Thermo Scientific, Wilmington, DE). Expression of macaque IRF3, IRF7, TRIM5α, and IFI16 vs. GAPDH was analyzed by one-step SYBR Green RT-qPCR (Kapa Biosystems, Wilmington, MA) according to the manufacturer’s instructions. Expression of IFNα vs. GAPDH was analyzed by two-step SYBR Green RT-qPCR (Kapa) as follows: cDNA was synthesized with the Superscript VILO cDNA synthesis kit, and Kapa SYBR Green RT-qPCR was performed. For all genes, primer concentrations were determined empirically and efficiency was determined prior to testing mRNA expression in samples. Data were analyzed by the ΔΔCt method. The cell control was GAPDH. The comparison control was sample from a single donor (same for all comparisons). The fold difference (2^-ΔΔCt^) is reported. Primer sequences are as follows. IRF3: F- 5’ CGCAGCCTCGAGTTTGAGAG 3’, R- 5’ ATGGTCCGGCCTACGATAGAA 3’; IRF7: F- 5’ ATGGGCAAGTGCAAGGTGTA 3’, R- 5’ ACCAGCTCTTGGAAGAAGACTC 3’; TRIM5α: F- 5’ GTGTGCCGGATCAGTTACCA 3’, R- 5’ GTCCCTCTTCTGGGCTCAAC 3’; IFI16: F- 5’ GAAGGCTGGAACGAAAGGGA 3’; R- 5’ GAAGGCTGGAACGAAAGGGA 3’; IFNα: F- 5’ AGCCTGTGTGATGCAGGAGATG 3’, R- 5’ GGAAGTATTTCCTCACGGCCAG 3’; GAPDH: F- 5’ GCTGAGTACGTCGTGGAGTC 3’, R- 5’ GGCGTTGCTGACGATCTTG 3’.

### Surface flow cytometry

Isolated rectal cells, PBMCs, and LNMCs were subjected to surface staining and flow cytometry as previously described (15, 83). Briefly, cells were incubated with the viability dye LIVE/DEAD Aqua (eBioscience), washed, and incubated with various antibody cocktails. Labeled cells were washed and fixed in 1% paraformaldehyde (PFA). All antibodies were purchased from Becton Dickenson (BD, Franklin Lakes, NJ) unless otherwise noted. Clone information is provided in parentheses for each antibody. Fluorescence minus one (FMO) controls were used throughout for gating. Data were acquired immediately after staining on an LSRII (BD) and analyzed with FlowJo software version 9.

Rectal cells were labeled with: CD3-AlexaFluor700 (SP34-2), CD4-V450 (L200), CD95-FITC (DX2), α4β7-APC (Non-human Primate Reagent Resource), CD69-APCH7 (FN50), CCR7-BV605 (Biolegend, San Diego, CA), CCR5-PE-Cy7 (NIH AIDS Reagent Program, labeled in-house), CCR6-PE (11-A9), and CD8-BUV395 (RPA-T8).

LNMCs and PBMCs were labeled with: CD3-APC-Cy7 (SP34-2), CD4-BUV395 (L200), CD200-BB515 (MRC OX-104), CD95-PE-Cy7 (DX2), CD127-BB700 (HIL-7R-M21), PD-1-PE-CF594 (EH12.1), CCR6-APC-R700 (11A9), CXCR3-BV605 (1C6), CD25-BV421 (M-A251), and CXCR5-PE (NHP, obtained from the Non-human Primate Reagent Resource).

### Intracellular cytokine flow cytometry

The frequencies of all cytokine-secreting T cells were monitored in jejunum. Isolated jejunum cells were stimulated with PMA (20 ng/mL)/ionomycin (0.5 μg/mL) vs. media for 1 hour at 37°C and then an additional 4 hours with Brefeldin A (10 μg/mL) and GolgiStop (BD) according to the manufacturer’s instructions. Stimulated cells were washed in Brilliant Staining Buffer (BD), stained with fixable viability stain (FVS)-575V (BD), and labeled with antibodies to surface markers (clones information is provided only if not provided above): CD3-APC-Cy7, CD4-BUV395, CD8-BUV496 (RPA-T8), and CD69-PerCP-Cy5.5 (FN50). Surface-labeled cells were then fixed in PFA overnight, permeabilized in BD Perm Wash Buffer, and labeled with antibodies to intracellular markers IFNγ-PE-Cy7 (4S.B3), IL17A-BV421 (N49-653), and IL22-APC (IL22JOP, eBioscience, San Diego, CA). Fully labeled cells were washed and the data were acquired immediately on an LSRII and analyzed with FlowJo software.

The frequencies of SIV-specific T cells were monitored in inguinal lymph nodes. Isolated LNMCs were incubated with SIV peptide pools covering gag and env in the presence of co-stimulatory αCD28 and αCD49d (BD, 10 μg/mL) on goat-α-mouse IgG F(ab)_2_-coated (KPL, Gaithersburg, MD) plates. Brefeldin A and GolgiStop were added after 1 hour and the cells stimulated for a further 5 hours. Stimulated cells were washed and stained with FVS-575V as above followed by surface staining, permeabilization, intracellular staining, and data acquisition and analysis as described above. Antibodies were the following: CD3-APC-Cy7, CD4-BUV395, CD8-BUV496, CD95-APC (DX2), CD69-PerCP-Cy5.5, CD40L-PE (TRAP1), TNFα-PE-CF594 (MAb11), IL2-APCR700 (MQ1-17H12), and IFNγ-PE-Cy7 (4S.B3).

### Co-cultures to detect replication competent SIV

5 x 10^4^ LNMCs were co-cultured with 10^5^ CEMx174 cells for 21 days. Culture media was exchanged every 3 days, and cultures were checked for signs of syncitia. On day 21, supernatants were collected for SIV p27 ELISA per the manufacturer’s instructions, and cells were collected for flow cytometry. The cells were stained for viability with Live/Dead Aqua stain, then labeled with antibody to CD4 (CD4-PE, L200), fixed and permeabilized with BD Fix/Perm, washed in BD Perm Wash, and incubated with Alexa647-conjugated antibody to SIV p27 (55-F12, provided by Mr. Trubey, NCI Frederick). Labeled cells were washed and the data acquired immediately on the LSRII and analyzed with FlowJo software.

### SIV env-specific antibody detection

Binding antibodies were detected by ELISA as follows. Heat-inactivated plasmas from baseline and the day of SIV challenge were incubated for 1 hour at 37°C on plates that had been coated with SIVmacA11 gp140 (NIH AIDS Reagent Program [ARP] Cat#2209 Lot16) in sodium bicarbonate buffer pH 9.6 overnight at 4°C and blocked 1 hour at 22°C in 2% bovine serum albumin (BSA)/PBS. Plates were washed in 1x ELISA plate wash (PerkinElmer, Waltham, MA) and anti-rhesus IgG-horseradish peroxidase (Non Human Primate Repository) was added for 1 hour at 37°C. Upon washing, the substrate was added for 0.5 hours at 22°C. The reaction was stopped with 1N hydrochloric acid and the optical density (OD) was read on an Emax Precision microplate reader with Softmax Pro software (Molecular Devices, San Jose, CA). Groups were compared at a plasma dilution of 1:2560.

Neutralizing antibodies were measured against lab-adapted SIVmac251. A stock of SIVmac251 was grown from stock obtained from the NIH ARP (Catalog #253) according to the protocol from the NIH ARP. Heat-inactivated plasmas from baseline and day of SIV challenge vs media were incubated with 50 TCID_50_ SIVmac251 in the presence of polybrene (4 μg/mL) on 5% BSA-coated plates for 1 hour at 37°C. Plasma/virus mixtures were then collected and added to 3 x 10^5^ CEMx174 cells for 7 days. On days 3 and 5 of culture, cells were fed with additional plasma or media. Sybr Green SIV RT-qPCR (Kapa) was used to measure SIV growth in the cultures, and SIV quantities in culture supernatant were determined by standard curve method. Percent neutralization was considered to be the SIV copy number in the presence of immune plasma subtracted from the SIV copy number in the presence of baseline plasma divided by the SIV copy number in the presence of baseline plasma.

### Statistics

Unless otherwise specified, the data were analyzed using the Kruskal Wallis test for unpaired samples with Dunns multiple comparison correction post-test. Significance level α was 0.05 throughout, but in order to identify trends, Wilcoxon Signed Rank test was performed for datasets with a Kruskal Wallis P<0.10, and all Kruskal Wallis P values less than 0.10 are shown. Everywhere multiple comparison correction was used, all comparisons were made except as noted in the figure legend. Mann Whitney test was used for binary comparisons (e.g. the effect of polyICLC on HSV-2 shedding). Spearman correlation coefficient was calculated to identify correlations between parameters (e.g. Tfh frequency and SIVΔNef plasma viral load). SIVΔNef take and SIV infection in macaques by treatment group was evaluated with two-sided Fisher’s exact test (control vs. treatment), and Chi Square test for trend (comparison of all groups).

## ACKNOWLEDGEMENTS

We would like to express our sincere thanks to Mr. Mac Trubey (AIDS and Cancer Virus Program, Frederick National Laboratory) for providing the SIV p27 antibody and staining method, and to Dr. Brandon Keele (AIDS and Cancer Virus Program, Frederick National Laboratory) for sequencing the *nef* gene of the plasma SIV in macaques with outlying viral loads to confirm the identity of the circulating virus. At the Population Council, we thank Drs. Elena Martinelli and Natalia Teleshova for helpful discussions and critical reading of the manuscript, Ms. Marlena Plagianos for discussions on statistical analysis, and Ms. Shimin Zhang for LSRII maintenance.

---

**Table 1. SIVΔNef uptake and SIV protection outcomes stratified by treatment group.** SIVΔNef take (ΔNEF+) is indicated as a fraction of LAV-challenged macaques that became infected with the vaccine. Within the ΔNEF+ macaque populations, the proportion protected at different levels is shown (“Protection from WT”). “Complete” protection is defined herein as no detectable SIV RNA in plasma above the limit of detection of the assay at any time. “Control” is defined as sustained SIV viremia <10^4^ copies/ml. “Any” protection refers to a group comprising both the completely protected and controller macaques. “No control” is defined as sustained viremia >10^4^ copies/ml. At each level of protection, the number of macaques in the group is also segmented by the time of SIV challenge post-LAV (i.e. 13 vs. 31 weeks). “MHC censored” indicates that animals with protective alleles Mamu A*01, B*08, and B*17 were excluded from analysis.

## SUPPORTING INFORMATION

**Figure S1. Plasma viral loads during the LAV and SIV phases of the study.** Plasma viral loads for SIVΔNef (filled symbols) and SIV (open symbols) are shown for each animal. Both viral loads were measured by PCR in *nef* or the region overlapping the *nef* deletion. Animals are grouped in columns according to level of protection from SIV and color coded according to treatment group as follows: HSV-2 (red), polyICLC (blue), HSV-2/polyICLC (purple), Untreated (black). MHC haplotypes Mamu*A01, *B08, and *B17 are indicated.

**Figure S2. MHC censored Figure 1.**

**Figure S3. MHC censored Figure 2.**

**Figure S4. MHC censored Figure 3.**

**Figure S5. Mucosal innate responses to low dose HSV-2 do not indicate immune dysregulation. (A)** Luminex data for representative immune mediators in clarified rectal swabs from 24 hours after the last HSV-2 and polyICLC treatment. **(B)** Luminex data for representative immune mediators in plasma from the same time point. **(C)** Surface phenotype of rectal CD4 T cells from 8 weeks post-SIVΔNef.

**Figure S6. MHC censored Figure 6.**

**Figure S7. MHC censored Figure 7.**

**Figure S8. MHC censored Figure 8.**

**Figure S9. MHC censored Figure 9.**

**Figure S10. SIV-specific CD4 T cell responses.** The frequency and phenotype of SIV gag- and env-specific CD4 T cells were analyzed by intracellular cytokine flow cytometry as in Figure 10 for CD8 T cells using LNMCs isolated 7 days post-SIV challenge from axillary lymph nodes of macaques challenged 31 weeks post-LAV. Data for CD4 T cells are shown. **(A)** Production of IFNγ (top), TNFα (middle), and IL2 (bottom) in response to SIV gag peptide pool vs media (No peptide) shown for a representative macaque. **(B)** Frequency of CD4 T cells expressing the indicated cytokine or surface marker in response to gag pool and env pool in each macaque shown by protection group. Each macaque is shown twice – once for gag response and once for env response. Colors and symbols are as in Figure 1. **(C)** Proportion of total CD4 T cell response attributable to each cytokine. Each macaque is shown twice, indicating the gag- and env-specific responses. **(D)** CD40L surface expression on CD8 T cells in response to PMA/ionomycin stimulation.

**Figure S11. MHC censored Figure 10.**

